## Supplementary Information for "PHEIGES, all-cell-free phage synthesis and selection from engineered genomes"

| Item | Title | Pages |
|---|---|---|
| Figure S1 | TXTL systems tests for remaining live *E. coli* cells | 2 |
| Figure S2 | Cell-free synthesis of deGFP | 3 |
| Figure S3 | Cell-free synthesis of T7 WT from its genome | 4 |
| Figure S4 | Construction of a plasmid using the DNA assembly method used for PHEIGES | 5 |
| Figure S5 | DNA electrophoresis gel of the four T7 WT PCR fragments | 6 |
| Figure S6 | PHEIGES to assemble the T7 WT phage | 7-8 |
| Figure S7 | PHEIGES to insert a *mcherry* cassette into the T7 WT | 9 |
| Figure S8 | Selection of T7-mCherry phages on plate reader | 10 |
| Figure S9 | PHEIGES with orthogonal oligonucleotides to insert a *mcherry* cassette into the T7 WT | 11 |
| Figure S10 | Infection kinetic assay of T7-mCherry phages on *E. coli* B cultures | 12 |
| Figure S11 | T7 phage information. | 13-14 |
| Figure S12 | List of *E. coli* strains used in this work and schematic of LPS | 15 |
| Figure S13 | Spotting of three T7 WT obtained with PHEIGES | 16 |
| Figure S14 | EOP of the T7 WT on all the *E. coli* strains used in this work | 17-18 |
| Figure S15 | In vitro genome ejection assay using purified LPS | 19 |
| Figure S16 | Co-expression experiment: link between genotype and phenotype. | 20-21 |
| Figure S17 | Dilution experiment: link between genotype and phenotype. | 22-23 |
| Figure S18 | Evaluation of randomness of tail fiber assembly in a cell-free environment. | 24-25 |
| Figure S19 | Generation of T7 phage variants with mutated tail fibers at 4 different rates of mutations | 26-27 |
| Figure S20 | Map of spotting assay of T7 phages with mutated tail fibers on *E. coli* B | 28 |
| Figure S21 | Spotting of T7-E0, T7-E1, T7-E2 and T7-E3 on all the *E. coli* strains used in this work | 29-30 |
| Figure S22 | Spotting of T7-E1, T7-E2 and T7-E3 on *rfaC*, *lpcA*, *rfaE*, *rfaD*, *rfaG* | 31 |
| Figure S23 | Spotting of T7 phage variants generated by PHEIGES on all the ReLPS *E. coli* strains | 32 |
| Figure S24 | Spotting of T7 phage variants generated by PHEIGES on strains with different LPS | 33 |
| Figure S25 | Mutation landscape for 12 T7 phage variants that infect strains *rfaD* and *rfaG* (ReLPS) | 34-35 |
| Figure S26 | Spotting of T7 phage variants with mutations only found in the tail genes *11* and *12* | 36 |
| Figure S27 | Spotting of T7 phage variants with mutations only found in the tail fiber gene *17* | 37 |
| Figure S28 | PHEIGES assembly of T7gp10-3xFLAG and T7gp17-3xFLAG | 38 |
| Figure S29 | Spotting of the T7-compilation phage on a lawn of the *E. coli rfaC* mutant strain | 39 |
| Figure S30 | In vitro genome ejection assay using purified LPS (T7-compilation) | 40 |
| Figure S31 | Infection kinetic assay of T7-ReLPS | 41 |
| Figure S32 | Spotting assay of five other phages expressed in TXTL | 42-43 |
| Figure S33 | PHEIGES of FelixO1 | 44 |

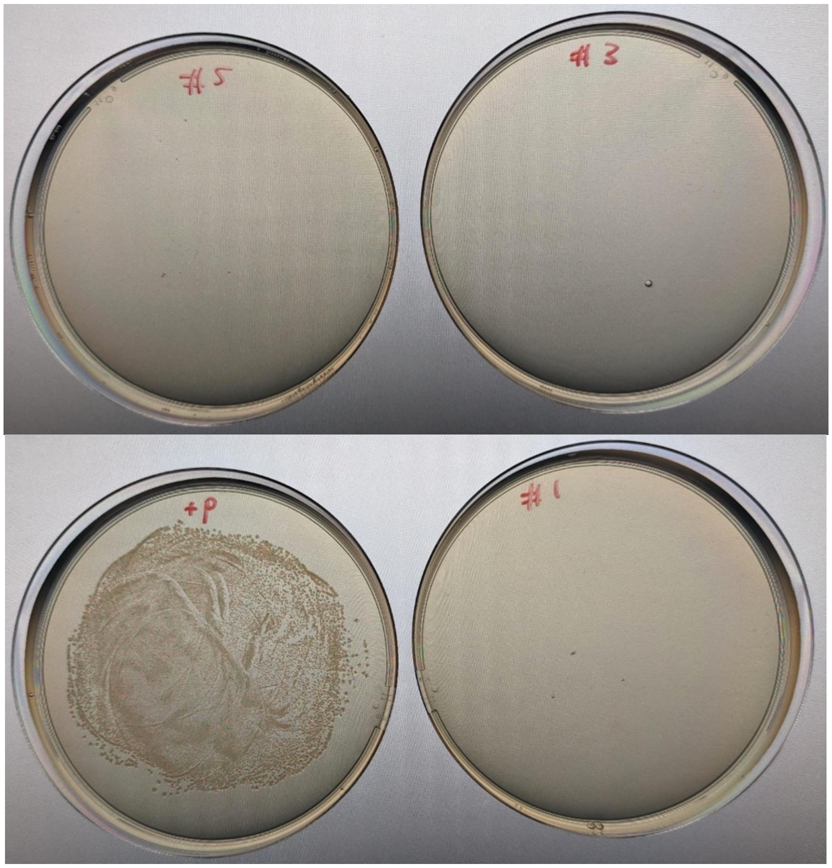

**Supplementary Figure 1.** TXTL systems were tested for remaining live *E. coli* cells. **Top row and bottom row left:** samples of three batches of TXTL systems (100 µl) were plated on Luria broth agar plates without antibiotics. The plates were incubated for 16 h at 37ºC. No colonies were visible. **Bottom right:** as a positive control, *E. coli* BL21 Δ*recBCD* Rosetta 2 cells were added to a TXTL sample to show that cells are viable in TXTL.

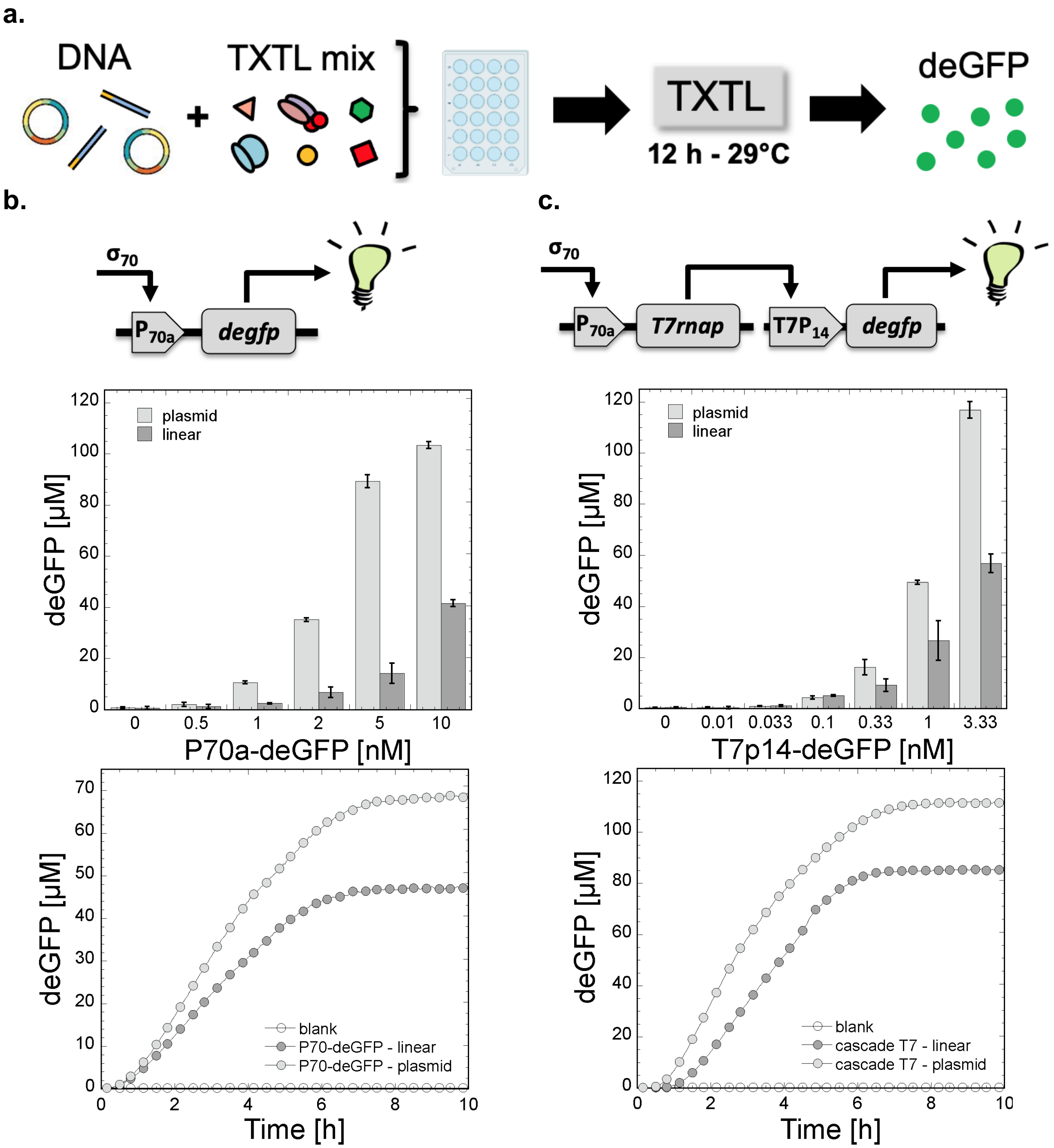

**Supplementary Figure 2.** Cell-free synthesis of deGFP. **a.** Schematic showing the TXTL workflow used in this work. Reactions were incubated on well plates. deGFP (a slightly shorter version of eGFP[1]) was synthesized either through the $P_{70a}$ promoter or through the T7 transcriptional activation cascade[1,2]. **b.** Endpoint cell-free synthesis of deGFP as a function of plasmid concentration ($P_{70a}$-*degfp*) and examples of kinetics. **c.** Endpoint cell-free synthesis of deGFP as a function of plasmid concentration ($P_{70a}$-*T7rnap* fixed at a concentration of 0.2 nM, T7p14-*degfp*) and examples of kinetics. Sequences of the plasmids are in **Table S4.**

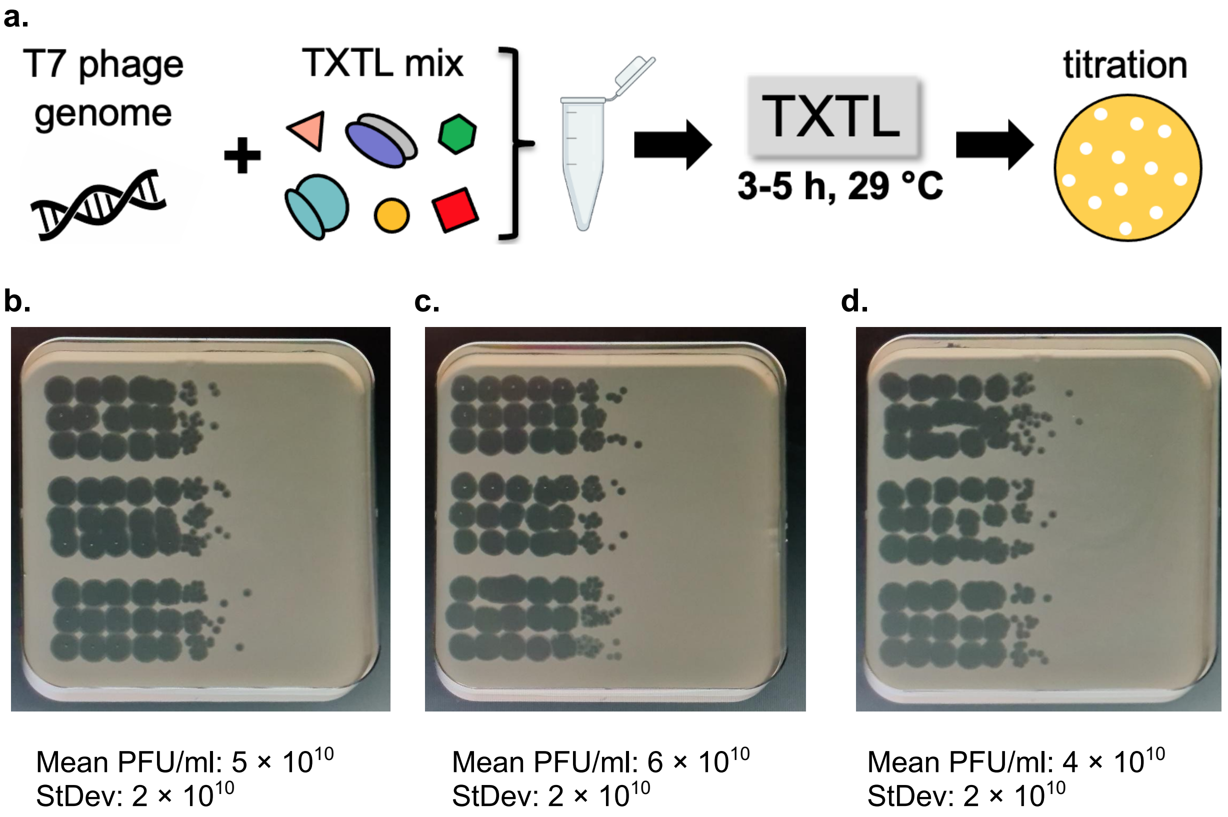

**Supplementary Figure 3.** Cell-free synthesis of T7 WT from its genome at 0.1 nM. **a.** Schematic of the experiment. The T7 WT was titrated by the spotting assay. The TXTL reactions were diluted in Luria broth by factors of ten. 3.5 µl of the dilution were spotted on a lawn of *E. coli* B cells from left to right. The first spot on the left corresponds to a dilution of 10 of the TXTL reaction. The last spots show single plaques, which enable determining the PFU/ml. The whole experiment was repeated three times and spotted three times (nine rows for each replicate). **b.** replicate 1. **c.** replicate 2. **d.** replicate 3.

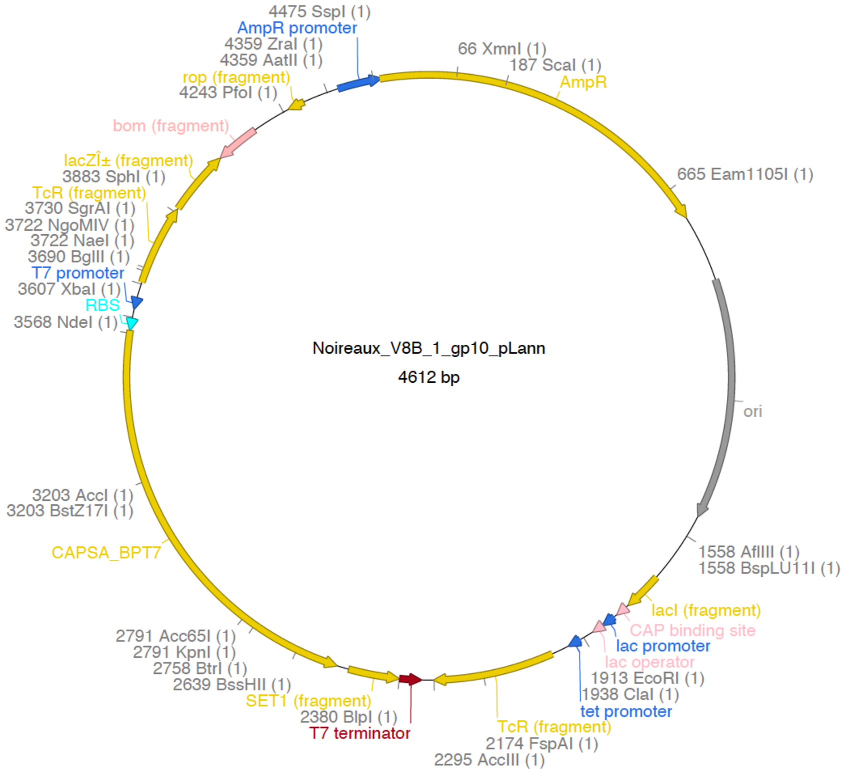

**Supplementary Figure 4.** Construction of a plasmid using the DNA assembly method used for PHEIGES. This plasmid was made from two PCR products: (i) one containing the ColE1 origin of replication and the ampicillin resistance gene, (ii) one containing the T7p14 promoter, the strong UTR1 RBS, the gene *10*, and the T7 terminator. The plasmid was fully sequenced and verified.

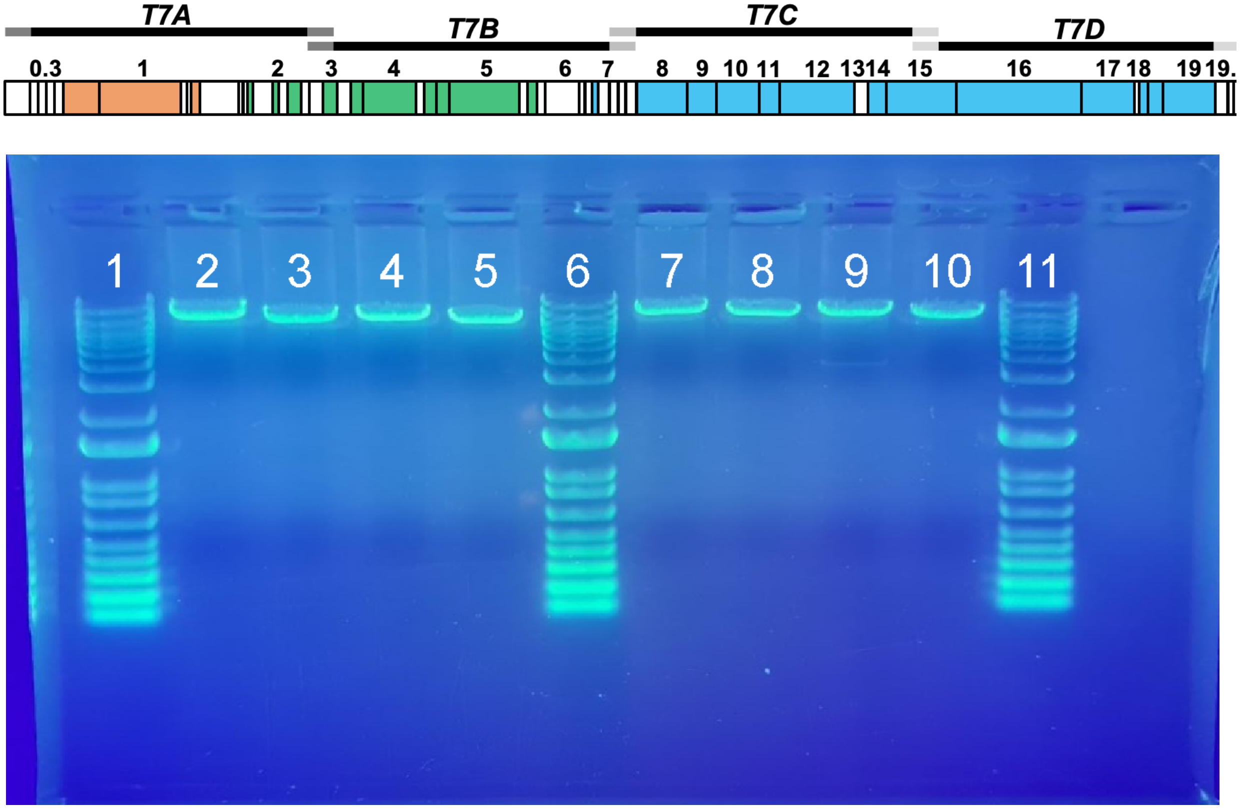

**Supplementary Figure 5.** DNA gel electrophoresis of the four T7 WT PCR fragments (T7A, T7B, T7C, T7D, as shown on the schematic of the T7 WT genome). The gel shows two sets of the fragments from two PCR runs. Lanes 1, 6, 11: 1 kb Plus DNA ladder (ThermoFisher). Lanes 2, 7: T7A (10.7 kbp). Lanes 3, 8: T7B (9.9 kbp). Lanes 4, 9: T7C (10.1 kbp). Lanes 5, 10: T7D (9.4 kbp).

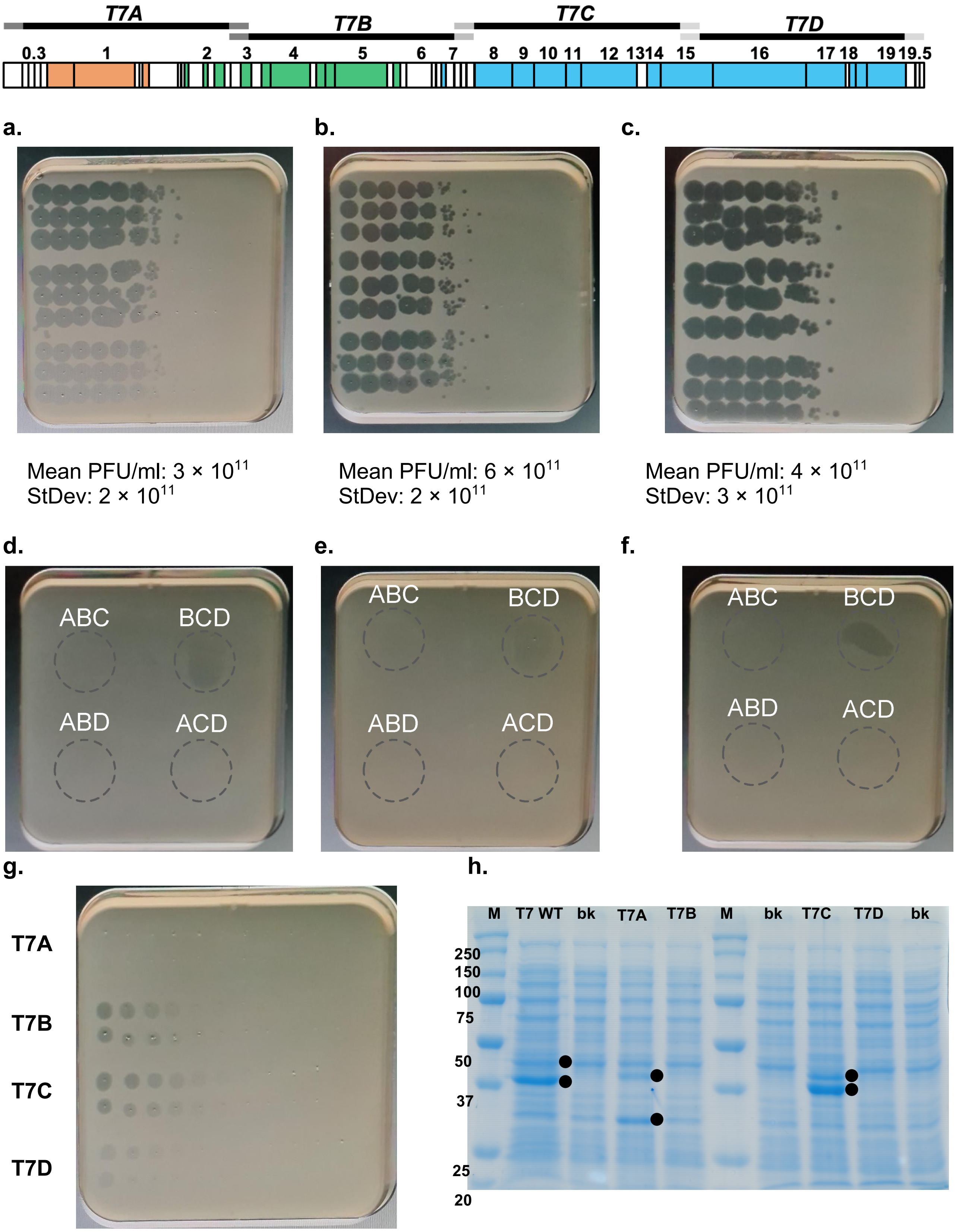
T7A
T7B
T7C
T7D
0.3
1
2
3
4
5
6
7
8
9
10
11
12
1314
15
16
17
18
1919.5
a.
b.
c.
Mean PFU/ml: 3 × 10^11
StDev: 2 × 10^11
Mean PFU/ml: 6 × 10^11
StDev: 2 × 10^11
Mean PFU/ml: 4 × 10^11
StDev: 3 × 10^11
d.
e.
f.
ABC
BCD
ABD
ACD
g.
T7A
T7B
T7C
T7D
h.
M
T7 WT
bk
T7A
T7B
M
bk
T7C
T7D
bk
250
150
100
75
50
37
25
20

**Supplementary Figure 6.** Using PHEIGES to assemble the T7 WT phage. The T7 WT phage was assembled from four DNA fragments T7A (10.7 kbp), T7B (9.9 kbp), T7C (10.1 kbp), T7D (9.4 kbp) to final equimolar mix at 0.1 nM. The experiment was repeated three times and spotted three times for each repeat. Three different batches of fragments were used. DNA fragments in repeats **a** and **b** were amplified from T7 genomic DNA (used at a concentration <1 fM), in repeat **c** the four DNA fragments were amplified by adding phages from a clarified phage lysate to the PCR mixed. The first spot on the left corresponds to a dilution of 10. Negative controls consisted of assembling the T7 WT from only three of the fragments. The experiment was repeated three times. Each repeat used a different batch of DNA fragments. DNA fragments in repeats **d** and **e** were amplified from T7 genomic DNA, in repeat **f** the four DNA fragments were amplified by adding phages from clarified phage lysate to the PCR mixed. No phages were detected. Some slight inhibition of *E. coli* growth was observed in some spots, especially for BCD. **g.** TXTL of four T7 genome parts in separate reactions and spotted on a lawn of *E. coli* B cells. Parts T7B, T7C and T7D show some inhibition activity. From left to right: each spot corresponds to a dilution (with water) by a factor of two of the TXTL reaction. The blank TXTL reaction (no DNA) does not show any inhibition on the lawn. **h.** SDS PAGE of the products of each genome part. The protein marker (M) shows the ladder masses in kDa. The blank reactions (bk) are TXTL reactions with no DNA added to them. The T7 WT shows two major bands at a size that corresponds to the capsid proteins 36.5 kDa and 41.8 kDa (products of genes *10A* and *10B*). The T7A part shows two proteins. One at a size of about 40 kDa, which corresponds to the protein kinase and/or the DNA ligase (products of gene *0.7* and gene *1.3*). The second band corresponds to a band of about 30 kDa, hypothetically corresponding to a single stranded DNA binding protein (product of gene *2.5*). The T7B part does not show any prominent bands. The T7C part shows two proteins, which corresponds to the capsid proteins 36.5 kDa and 41.8 kDa (products of genes *10A* and *10B*). The T7D part does not show any prominent bands. For T7B, T7C and T7D, proteins are synthesized without the T7 RNA polymerase being produced.

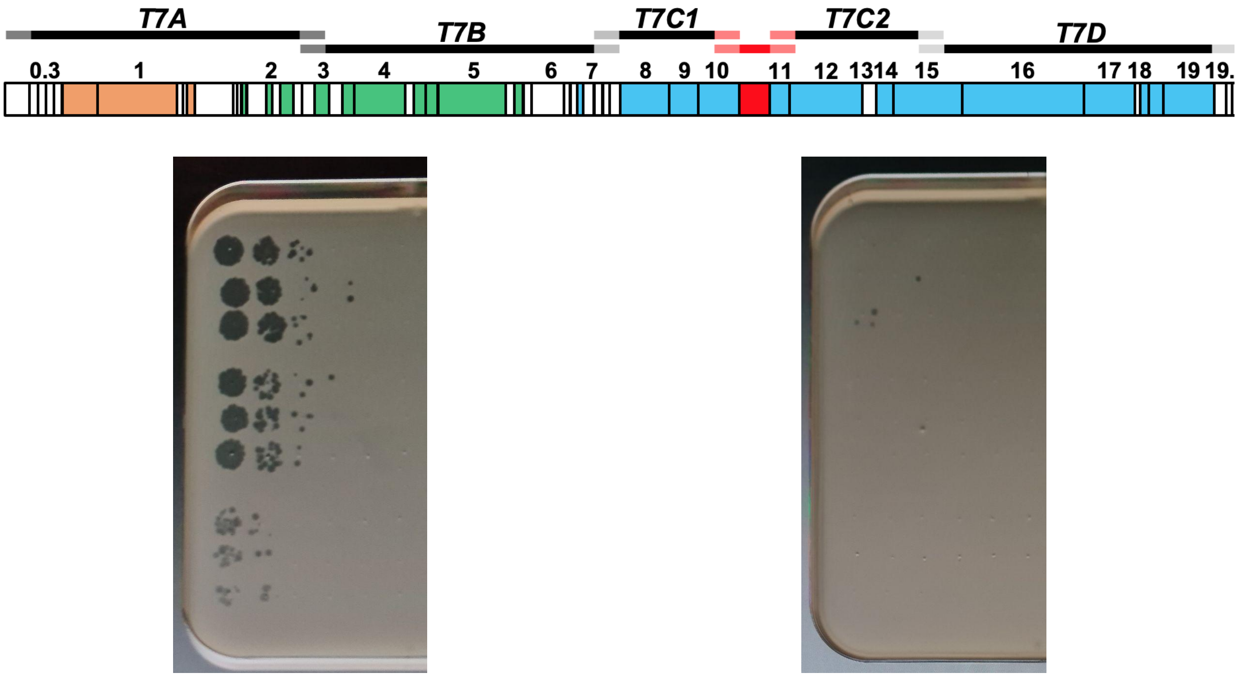

Mean PFU/ml: $9 \times 10^6$
StDev: $7 \times 10^6$

Mean PFU/ml: $2 \times 10^3$
StDev: $2 \times 10^3$

**Supplementary Figure 7.** Using PHEIGES to insert a *mcherry* gene cassette into the T7 WT after gene *10A*, before the T7 terminator of gene *10A*. This experiment was carried out using oligonucleotides not designed to be specifically orthogonal. **Left:** assembly of the five DNA fragments including the *mcherry* cassette. The first spot on the left corresponds to a dilution of 10 of the TXTL reaction after PHEIGES. **Right:** assembly of the four DNA fragments without the *mcherry* cassette. Although resulting in a PFU/ml one thousand time smaller, a significant number of phages are assembled without the *mcherry* gene. After this result, we demonstrated that using orthogonal oligonucleotides for the DNA assembly eradicates leaky synthesis of phages, thus creating a leak-free PHEIGES workflow.

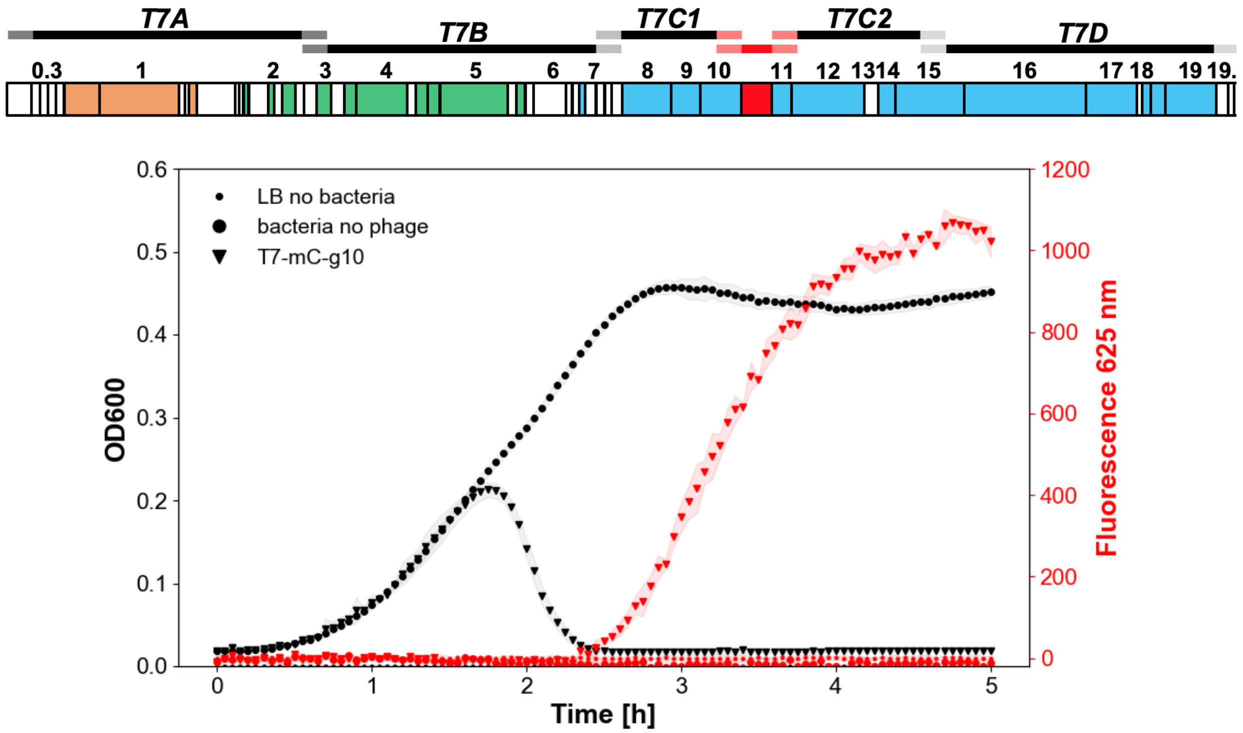

**Supplementary Figure 8.** Selection of T7-mCherry phages on plate reader. PHEIGES was used without orthogonal oligonucleotides to insert a *mcherry* gene cassette into the T7 WT after gene *10A*, before the T7 terminator of gene *10A*, as shown in **Fig. S7**. A serial dilution of the TXTL reaction expressing and synthesizing the T7-mCherry phages was added to *E. coli* B cultures to isolate phages carrying the *mcherry* gene cassette. With a concentration one thousand greater than T7 WT, T7-mCherry phages were isolated by taking the last dilution that shows mCherry fluorescence. The kinetics of the OD600 and fluorescence shows one T7-mCherry phage variant after selection.

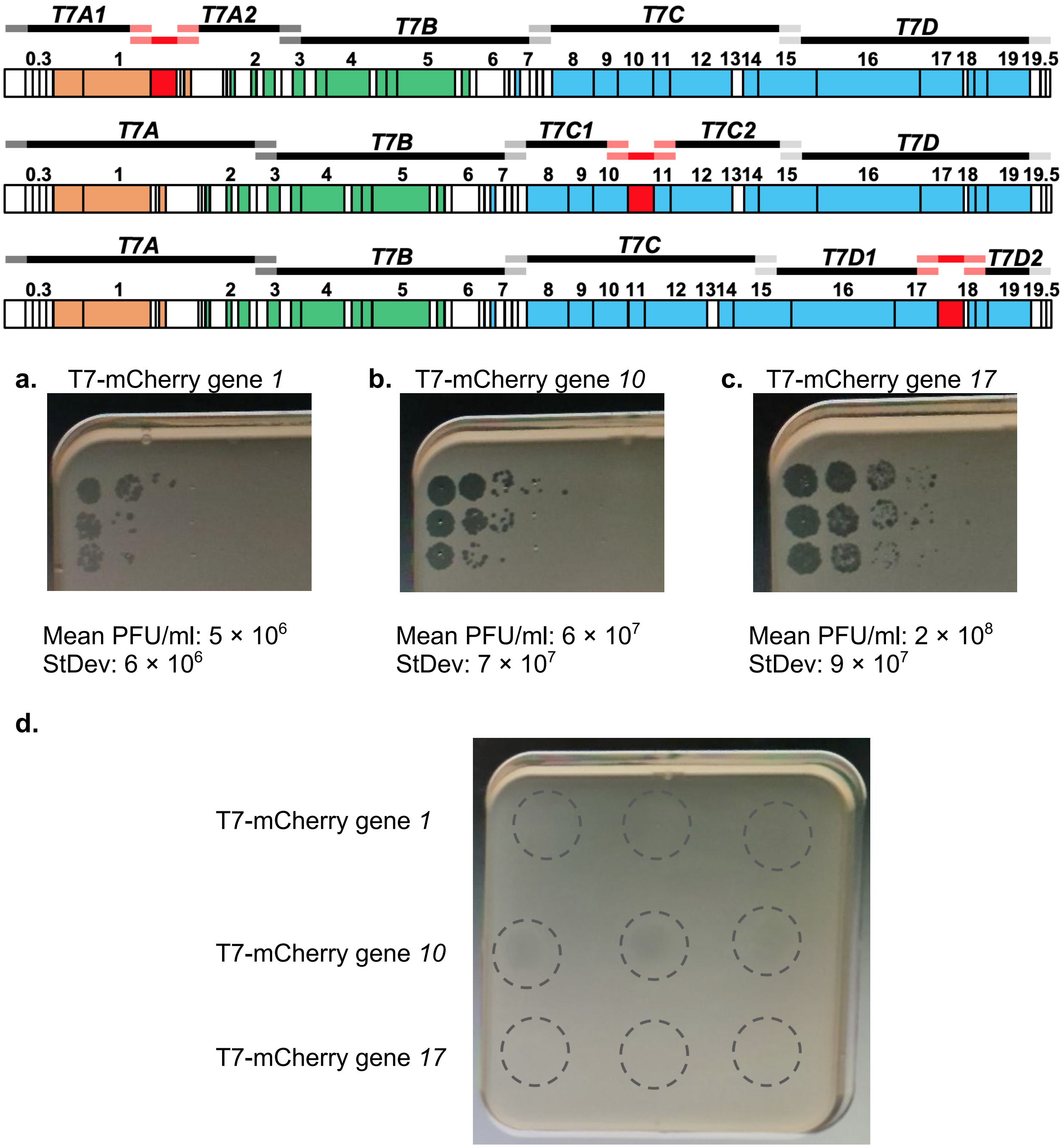

**Supplementary Figure 9.** Using PHEIGES with orthogonal oligonucleotides to insert a *mcherry* gene cassette into the T7 WT after gene *1*, gene *10A*, and gene *17*. **a.** Insertion after gene *1*. **b.** Insertion after gene *10A*. **c.** Insertion after gene *17*. The PFU/ml indicate the efficacy of the insertion, which depends up to a hundred-fold on the loci in this experiment. This experiment was replicated three times (three rows). The first spot on the left corresponds to a dilution of 10. **d.** The controls consisted of the same experiment without including the *mcherry* gene cassette in the DNA assembly. Negative controls were repeated three times. No phages were synthesized.

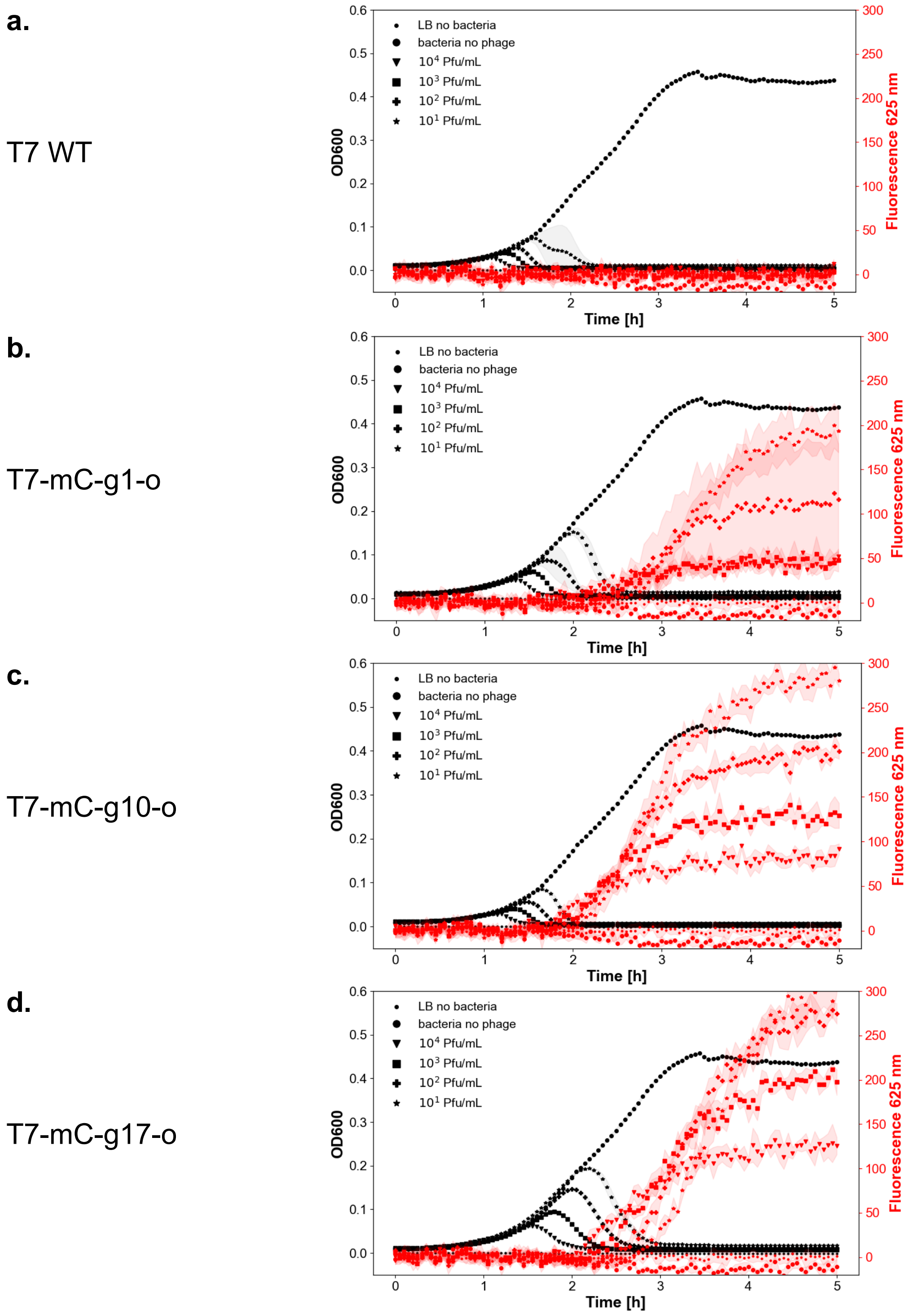

**Supplementary Figure 10.** Infection kinetic assay of T7-mCherry phages (assembled with orthogonal oligonucleotides) on *E. coli* B cultures. The left axis shows the OD600 (dark curves), the right axis shows the fluorescence at 625 nm (red curves). **a.** T7 WT. **b.** T7-mC-g1-o. **c.** T7-mC-g10-o. **d.** T7-mC-g17-o.

**a.**

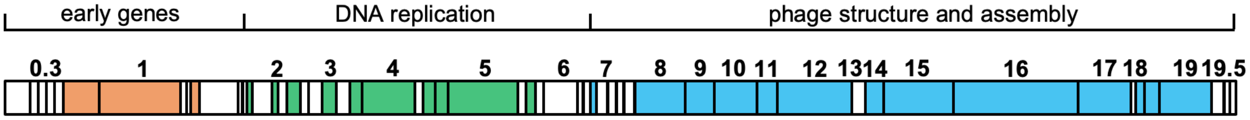

**b.**

| Gene name | Position | Function | Structural | Size in kDa |
|---|---|---|---|---|
| *0.3* | 925-1278 | inhibits EcoB/K | No | 13.8 |
| *0.4* | 1278-1433 | inhibit host cell division | No | 5.7 |
| *0.5* | 1496-1639 | unknown | unknown | 4.7 |
| *0.6A and B* | 1636-1797 | unknown | unknown | 6.2 and 13.2 |
| *0.7* | 2021-3100 | protein kinase | No | 41.1 |
| *1* | 3171-5822 | T7 RNA polymerase | No | 98.8 |
| *1.1* | 6007-6135 | unknown | unknown | 5.2 |
| *1.2* | 6137-6394 | inhibits dGTP triphosphohydrolase | No | 10.2 |
| *1.3* | 6475-7554 | DNA ligase | No | 41.1 |
| *1.4* | 7608-7763 | unknown | unknown | 5.45 |
| *1.5* | 7791-7880 | unknown | unknown | 3.1 |
| *1.6* | 7906-8166 | unknown | unknown | 9.9 |
| *1.7* | 8166-8756 | nucleotide kinase | No | 22.2 |
| *1.8* | 8749-8895 | unknown | unknown | 5.8 |
| *2* | 8898-9092 | inhibits *E. coli* RNA polymerase | No | 7.18 |
| *2.5* | 9158-9856 | single-stranded DNA binding protein | No | 25.7 |
| *2.8* | 9857-10276 | unknown | unknown | 15.6 |
| *3* | 10257-10706 | endonuclease | No | 17.2 |
| *3.5* | 10706-11161 | lysozyme, inhibits T7 RNA polymerase | No | 16.98 |
| *3.8* | 11125-11590 | unknown | unknown | 3.8 |
| *4* | 11565-13265 | primase helicase | No | 62.6 |
| *4.1* | 11635-11757 | unknown | unknown | 4.4 |
| *4.2* | 12988-13326 | unknown | unknown | 12.65 |
| *4.3* | 13352-13564 | unknown | unknown | 7.9 |
| *4.5* | 13584-13853 | inhibitor of toxin/antitoxin system | No | 10.1 |
| *4.7* | 13927-14334 | unknown | unknown | 15.2 |
| *5* | 14353-16467 | DNA polymerase | No | 79.7 |
| *5.3* | 16483-16839 | unknown | unknown | 13.07 |
| *5.5* | 16851-17150 | permit growth on lambda lysogen | No | 11.2 and 18.7 |
| *5.7* | 17150-17359 | unknown | unknown | 7.4 |
| *5.9* | 17359-17517 | inhibits host recBCD | No | 6.04 |
| *6* | 17504-18406 | exonuclease | No | 34.5 |
| *6.3* | 18394-18507 | unknown | unknown | 4.09 |
| *6.5* | 18605-18859 | unknown | unknown | 9.47 |
| *6.7* | 18864-19130 | head protein | Yes | 9.34 |
| *7* | 19130-19531 | host range | No | 15.4 |
| *7.3* | 19535-19834 | tail protein | Yes | 10.07 |
| *7.7* | 19848-20240 | unknown | unknown | 14.7 |
| *8* | 20240-21850 | head-tail connector protein | Yes | 59.1 |
| *9* | 21950-22873 | scaffolding protein | Yes | 33.9 |
| *10A and B* | 22967-24162 | major and minor capsid protein | Yes | 36.5 and 41.8 |
| *11* | 24228-24818 | tail protein | Yes | 22.29 |
| *12* | 24842-27226 | tail protein | Yes | 89.4 |
| *13* | 27307-27723 | chaperone like activity | No | 15.85 |
| *14* | 27728-28318 | internal virion protein | Yes | 20.97 |
| *15* | 28325-30568 | internal virion protein | Yes | 84.34 |

| | | | | |
|---|---|---|---|---|
| *16* | 30595-34551 | internal virion protein | Yes | 143.84 |
| *17* | 34646-36285 | tail fiber protein | Yes | 61.57 |
| *17.5* | 36344-36547 | lysis protein | No | 7.39 |
| *18* | 36553-36822 | DNA maturation protein | unknown | 10.15 |
| *18.5* | 36917-37348 | lysis protein | No | 16.24 |
| *18.7* | 37032-37283 | unknown | unknown | 9.32 |
| *19* | 37370-39130 | DNA maturation protein | unknown | 66.26 |
| *19.2* | 38016-38273 | unknown | unknown | 9.39 |
| *19.3* | 38553-38726 | unknown | unknown | 6.56 |
| *19.5* | 39389-39538 | unknown | unknown | 5.43 |

**Supplementary Figure 11.** T7 phage information. **a.** Schematic of the genome adapted from *Bacteriophage protein-protein interactions*[3] with some of the genes annotated. **b.** List of the genes and positions, with their functions when known and the size of the proteins (from GenBank V01146.1).

| strain | LPS type | gene name | provider |
|---|---|---|---|
| *E. coli* B | *E. coli* B LPS – analogue to RbLPS | | ATCC 11303 |
| Strain Seattle 1946 | smooth (O6) | | ATCC 25922 |
| ClearColi BL21(DE3) | IVa | | Lucigen |
| *rfaC* | Re | *waaC* | Keio collection |
| *lpcA* | Re | *gmhA* | Keio collection |
| *rfaE* | Re | *gmhE* | Keio collection |
| *rfaD* | Re | *gmhD* | Keio collection |
| *gmhB* | Re partial | *gmhB* | Keio collection |
| *rfaF* | Rd2 | *waaF* | Keio collection |
| *rfaG* | Rd1 | *waaG* | Keio collection |
| *galU* | Rd1 (leaky) | *galU* | Keio collection |
| *rfaI* | Rc | *waaO / waaI* | Keio collection |
| *rfaJ* | Rb | *waaR* | Keio collection |
| *rfaB* | LPS side-chain additions/modifications | *waaB* | Keio collection |
| *rfaL* | LPS side-chain additions/modifications | *waaL* | Keio collection |
| *rfaP* | LPS side-chain additions/modifications | *waaP* | Keio collection |
| *rfaQ* | LPS side-chain additions/modifications | *waaQ* | Keio collection |
| *rfaS* | LPS side-chain additions/modifications | *waaS* | Keio collection |
| *rfaY* | LPS side-chain additions/modifications | *waaY* | Keio collection |
| *rfaZ* | LPS side-chain additions/modifications | *waaZ* | Keio collection |

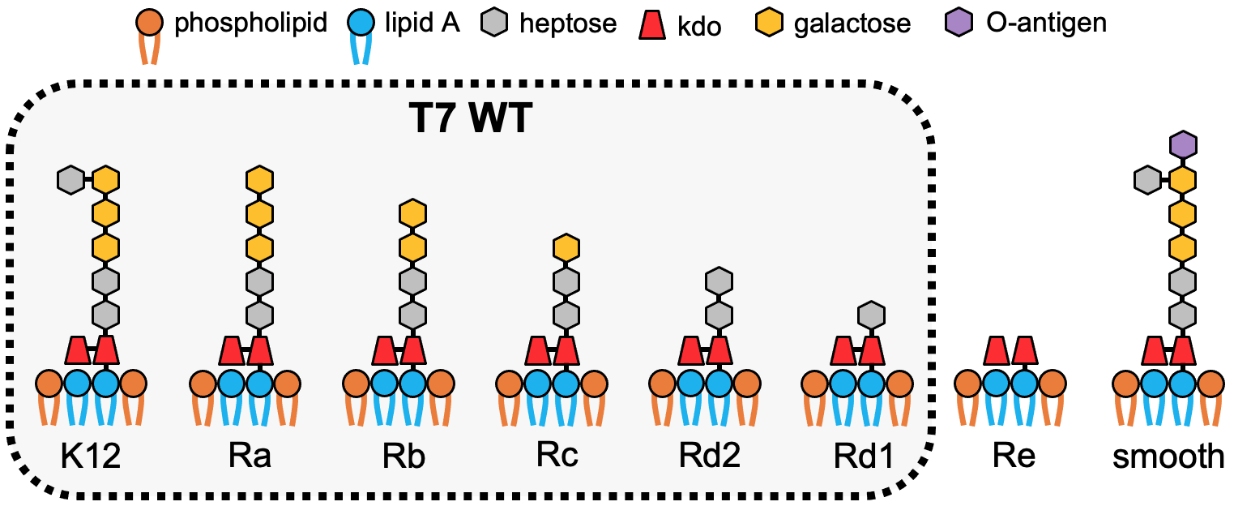

**Supplementary Figure 12. Top:** list of *E. coli* strains used in this work. This table is also in the Supplementary Tables file. **Bottom:** schematic of the LPS and the ones recognized by T7 WT (boxed). Note that *E. coli* B is a RbLPS type[4].

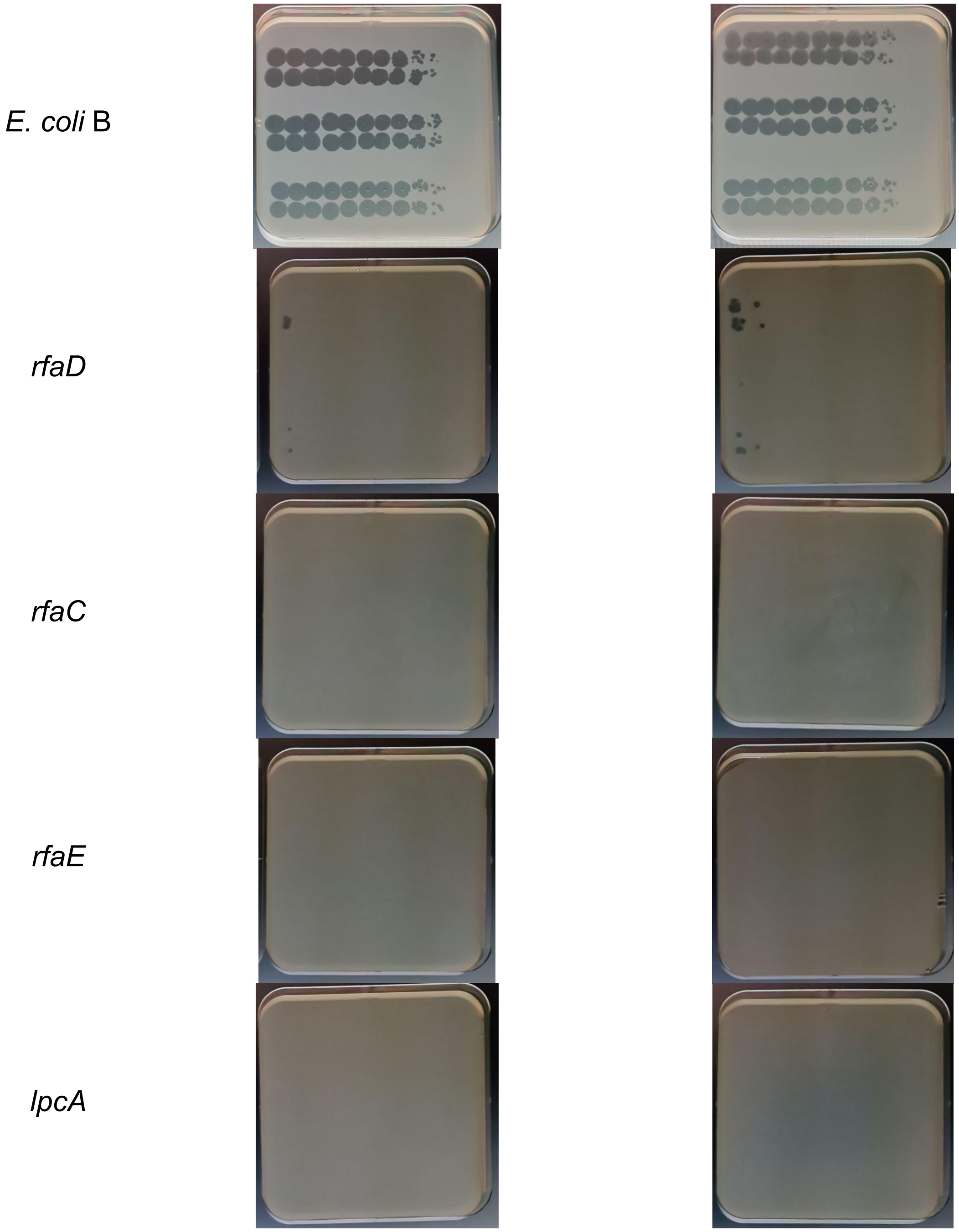

**Supplementary Figure 13.** Spotting assay of three T7 WT lysates obtained from TXTL reactions using the T7 WT genome (Boca Scientific), the genome assembled from four fragments using PHEIGES, and from T7-E0. Isolated plaques were picked and amplified in *E. coli* B and clarified lysates were diluted to $10^8$ PFU/ml. The genomes of the three phages were fully sequenced. Each phage lysate was spotted twice and each on *E. coli* B, and the mutant strains *rfaC*, *rfaE*, *lpcA* and *rfaD*. Plaques are detected in all lysates on *rfaD* but not on *rfaC*, *lpcA* and *rfaE*. Spotting was done from left to right. The first spot corresponds to no dilution, serial 5-fold dilutions for the other. Left and right columns are replicates.

**a.**

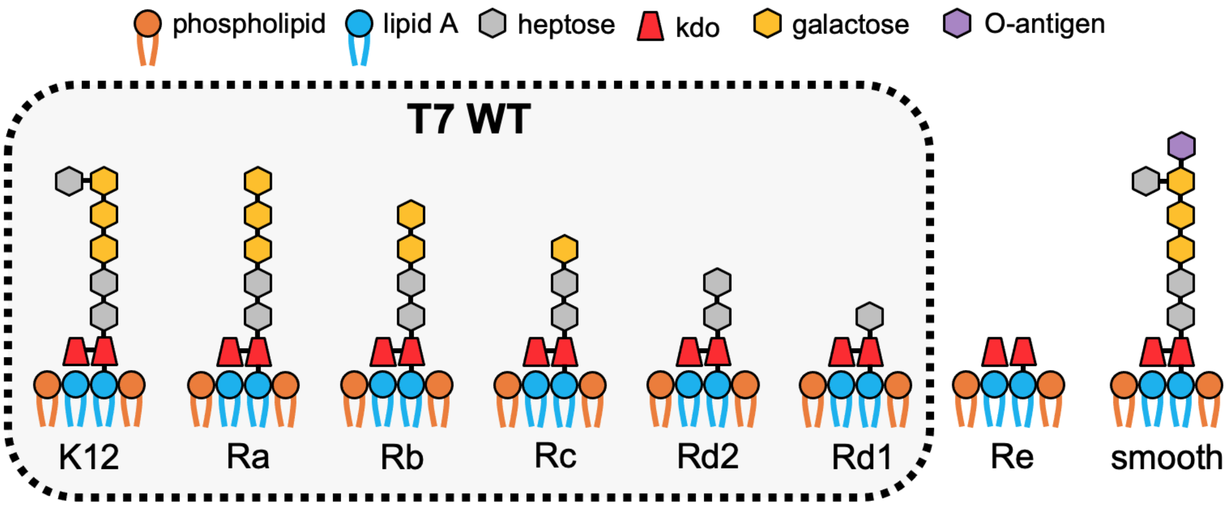

| strains | LPS type | EOP |
|---|---|---|
| | | T7-E0 |
| *BW25113* | *BW25113* K12 LPS | 1 |
| *E. coli* B | *E. coli* B LPS | 1 |
| Strain Seattle 1946 | smooth (O6) | no plaque |
| ClearColi BL21(DE3) | IVa | no plaque |
| *rfaC* | Re | no plaque ($<10^{-5}$) |
| *lpcA* | Re | no plaque ($<10^{-5}$) |
| *rfaE* | Re | no plaque ($<10^{-5}$) |
| *rfaD* | Re | 0.0004 |
| *gmhB* | Re partial | 0.8 |
| *rfaF* | Rd2 | 0.4 |
| *rfaG* | Rd1 | 0.04 |
| *galU* | Rd1 (leaky) | 0.6 |
| *rfaI* | Rc | 0.8 |
| *rfaJ* | Rb | 0.1 |
| *rfaB* | LPS side-chain additions/modifications | 0.8 |
| *rfaL* | LPS side-chain additions/modifications | 0.8 |
| *rfaP* | LPS side-chain additions/modifications | 0.6 |
| *rfaQ* | LPS side-chain additions/modifications | 0.6 |
| *rfaS* | LPS side-chain additions/modifications | 0.9 |
| *rfaY* | LPS side-chain additions/modifications | 0.8 |
| *rfaZ* | LPS side-chain additions/modifications | 0.9 |

**b**.

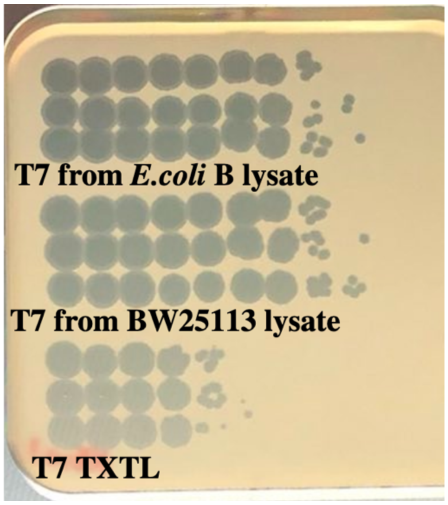

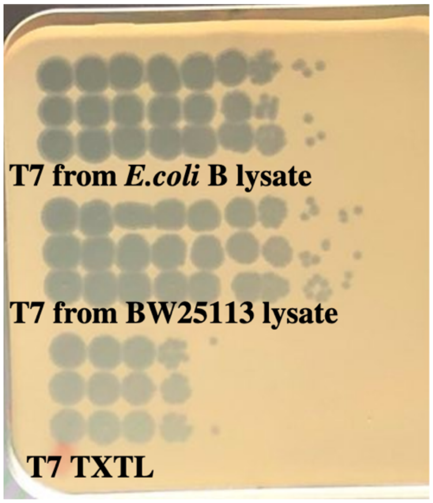

**Titers on *BW25113*:**

T7 *E. coli* B lysate: $4.5 \pm 1 \times 10^{11}$ PFU/ml
T7 *BW25113* lysate: $5.5 \pm 0.8 \times 10^{11}$ PFU/ml
T7 TXTL: $3.0 \pm 0.8 \times 10^{8}$ PFU/ml

**Titers on *E. coli* B:**

T7 *E. coli* B lysate: $1.8 \pm 1 \times 10^{11}$ PFU/ml
T7 *BW25113* lysate: $5.8 \pm 0.9 \times 10^{9}$ PFU/ml
T7 TXTL: $1 \pm 0.5 \times 10^{8}$ PFU/ml

**Supplementary Figure 14. a**. EOP of the T7 WT on all the *E. coli* strains used in this work. **b**. T7 lysate from *E. coli* B, KEIO parent strain *BW25113* and TXTL expressed. Similar titers are observed in all conditions. This suggests that *E. coli* B and K12 derivative strains modification-restriction systems do not affect T7 EOP and that T7 interacts with *E. coli* B and *E. coli* BW25113 in the same way.

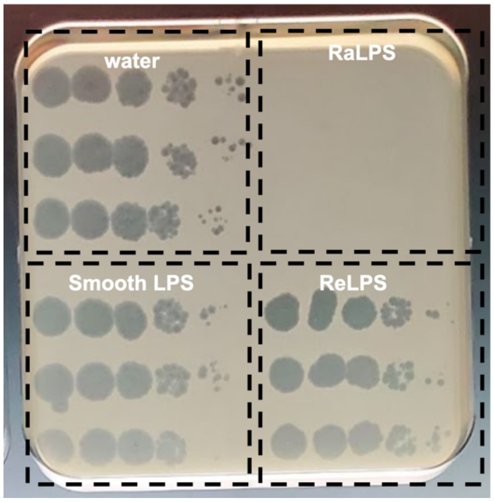

T7 WT phage spotting on a lawn of *E. coli* B after incubation with:

| | mean PFU/ml | StDev |
|---|---|---|
| + water | $2 \times 10^8$ | $5 \times 10^7$ |
| + smooth LPS | $2 \times 10^8$ | $1 \times 10^8$ |
| + RaLPS | $<10^3$ | - |
| + ReLPS | $2 \times 10^8$ | $1 \times 10^8$ |

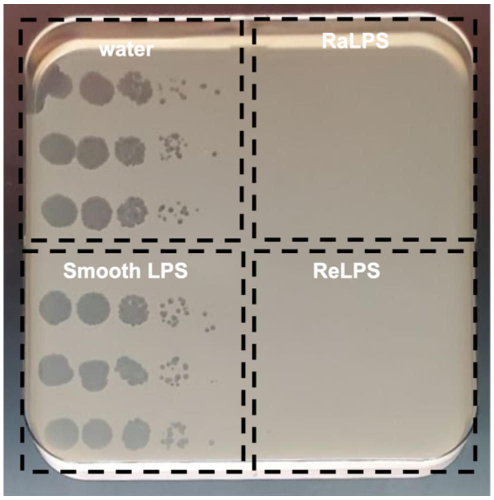

T7-ReLPS phage spotting on a lawn of *E. coli* B after incubation with:

| | mean PFU/ml | StDev |
|---|---|---|
| + water | $4 \times 10^7$ | $3 \times 10^7$ |
| + smooth LPS | $5 \times 10^7$ | $3 \times 10^7$ |
| + RaLPS | $<10^3$ | - |
| + ReLPS | $<10^3$ | - |

**Supplementary Figure 15.** In vitro genome ejection assay using purified LPS. Spotting (on a lawn of *E. coli* B) of serial 10-fold dilutions of the T7 WT and a T7-rfaD-1 phages. The two phages at ~$10^9$ PFU/mL were pre-incubated with purified LPS at 0.2 mg/mL at 37 °C for 2 hours. RaLPS is the full size LPS. In the case of T7 WT, only the RaLPS induces ejection of the genomes during the pre-incubation, resulting in no plaques. In the cases of T7-ReLPS both the RaLPS and the ReLPS induce ejection of the genome during the pre-incubation resulting in no plaques. The first spot on the left corresponds to no dilution. The titer loss of T7-rfaD-s compared to T7 WT in presence of ReLPS shows that T7-rfaD-1 is at least ~10 000 times (T7WT-ReLPS/T7WT-water)/(T7ReLPS-ReLPS/T7ReLPS-water) more selective to ReLPS in vitro than T7 WT at 0.2 mg/mL ReLPS 37 °C, 2h. Each condition corresponds to tree individual phage/LPS mixture spotted once.

**a.** ***E. coli* B**

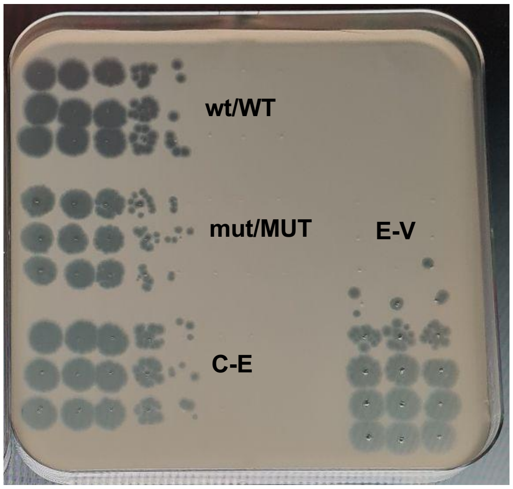

***rfaC***

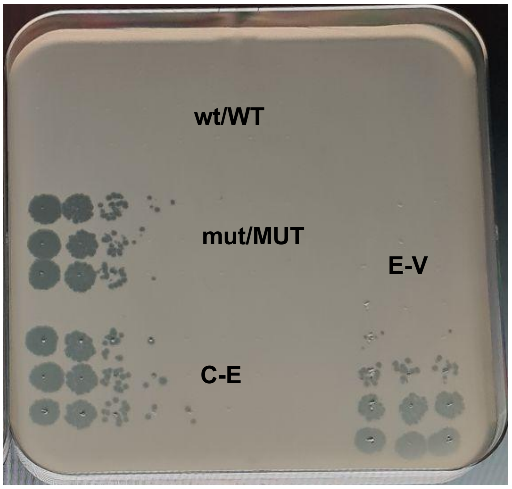

**Titers on *E. coli* B:**

wt/WT: $4.2 \pm 0.8 \times 10^{9}$ PFU/ml
mut/MUT: $2.9 \pm 0.1 \times 10^{9}$ PFU/ml
C-E: $4.9 \pm 0.4 \times 10^{9}$ PFU/ml
E-V: $3.9 \pm 0.4 \times 10^{9}$ PFU/ml

**Titers on *rfaC*:**

wt/WT: $<10^{5}$ PFU/ml, no plaque detected
mut/MUT: $6.3 \pm 0.9 \times 10^{8}$ PFU/ml
C-E: $4.4 \pm 0.4 \times 10^{8}$ PFU/ml
E-V: $4.0 \pm 0.7 \times 10^{8}$ PFU/ml

**b.** ***E. coli* B**

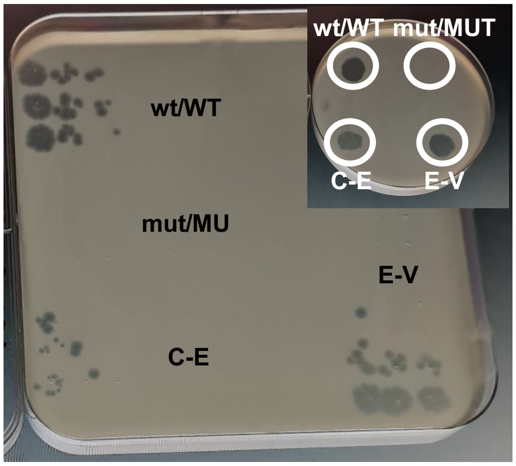

***rfaC***

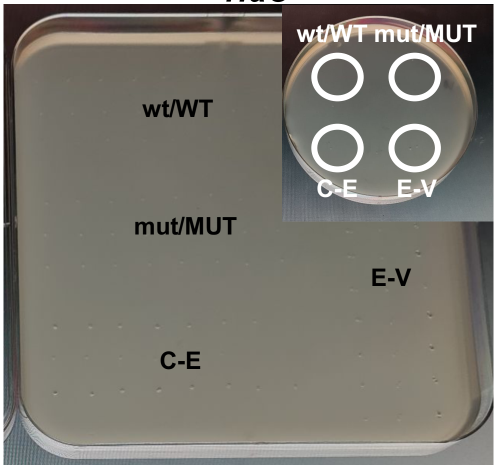

**Titers on *E. coli B*:**

wt/WT: $2.3 \pm 0.4 \times 10^{7}$ PFU/ml
mut/MUT: 0 PFU/ml, no plaque detected
C-E: $1.7 \pm 0.5 \times 10^{7}$ PFU/ml
E-V: $1.6 \pm 0.2 \times 10^{8}$ PFU/ml

**Titers on *rfaC*:**

wt/WT: 0 PFU/ml, no plaque detected
mut/MUT: 0 PFU/ml, no plaque detected
C-E: 0 PFU/ml, no plaque detected
E-V: 0 PFU/ml, no plaque detected

**c.**

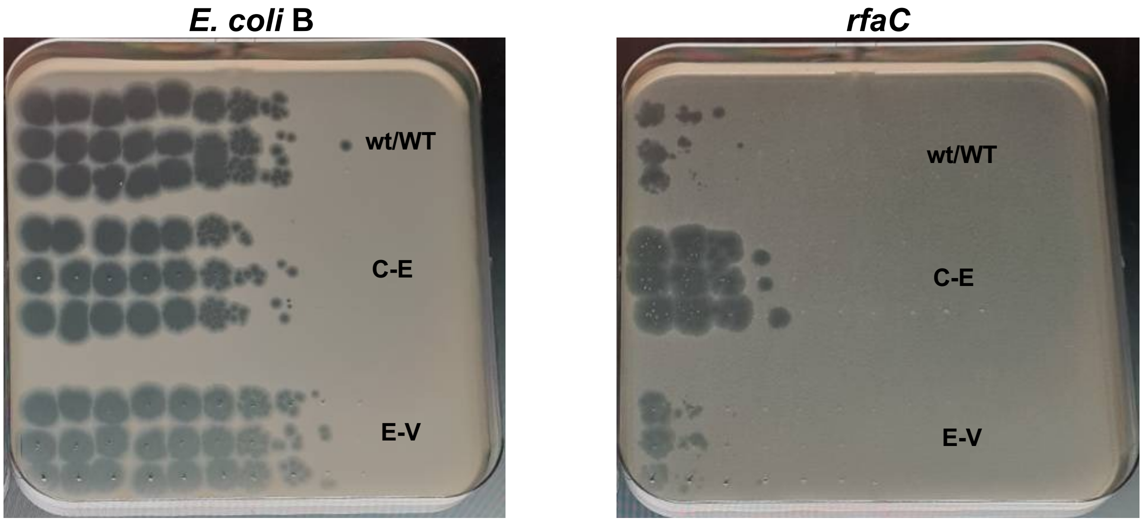

**Titers on *E. coli* B:**
wt/WT: $1.2 \pm 1 \times 10^{11}$ PFU/ml
C-E: $1.7 \pm 0.4 \times 10^{10}$ PFU/ml
E-V: $2.1 \pm 0.5 \times 10^{11}$ PFU/ml

**Titers on *rfaC*:**
wt/WT: $6.7 \pm 3 \times 10^{4}$ PFU/ml
C-E: $2.5 \pm 0.1 \times 10^{6}$ PFU/ml
E-V: $1.0 \pm 0.3 \times 10^{5}$ PFU/ml

**Supplementary Figure 16.** Co-expression experiment: link between genotype and phenotype. **a.** Spotting assay of the four phage solutions on strains *E. coli* B and *rfaC* expressed at final 0.1 nM of DNA. C-E: co-expression. E-V: equal volume. The first spot corresponds to a dilution of 100. **b.** Ejection assay with 0.4 mg/ml of ReLPS 37 °C overnight (stringent conditions), applied to the four phage solutions. Spotting assay of the four phage solutions on strains *E. coli* B and *rfaC* after ejection. The first spot corresponds to a dilution of 1000 of the original TXTL reaction. Inset: 25 µl of each of the four phage solutions were spotted without any dilution. No plaques were detected after ejection on the *rfaC* strain. **c.** Amplification in *E. coli* B and spotting on *E. coli* B and *rfaC*. The first spot corresponds to no dilution. No lysis occurred for the ejection for M/M only. Each condition is three reactions spotted once.

**a.** *E. coli* **B** *rfaC*

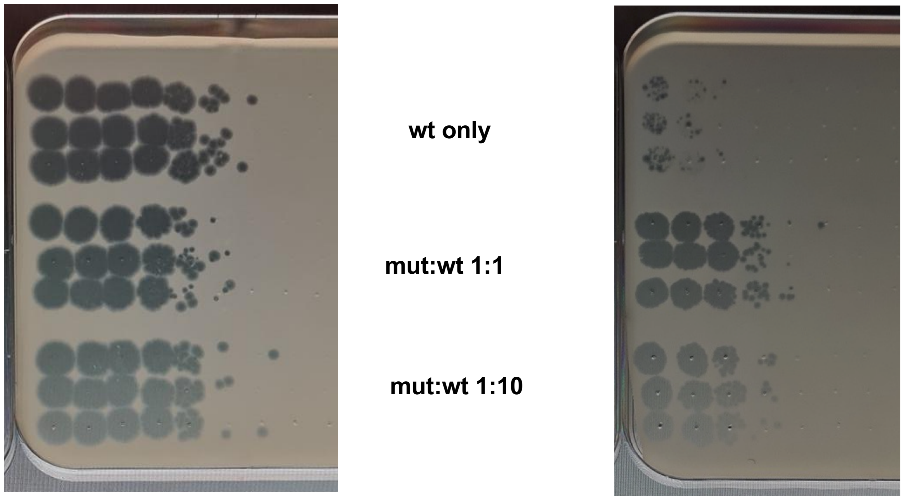

**Titers on *E. coli* B:**
wt only: $1.7 \pm 0.3 \times 10^{9}$ PFU/ml
mut:wt: 1:1 $5.0 \pm 1.4 \times 10^{8}$ PFU/ml
mut:wt: 1:10 $3.3 \pm 1.4 \times 10^{8}$ PFU/ml

**Titers on *rfaC*:**
wt only: $7.7 \pm 1.2 \times 10^{5}$ PFU/ml
mut:wt: $2.3 \pm 0.7 \times 10^{8}$ PFU/ml
mut:wt: $5.0 \pm 0.9 \times 10^{7}$ PFU/ml

**b.** *E. coli* **B** *rfaC*

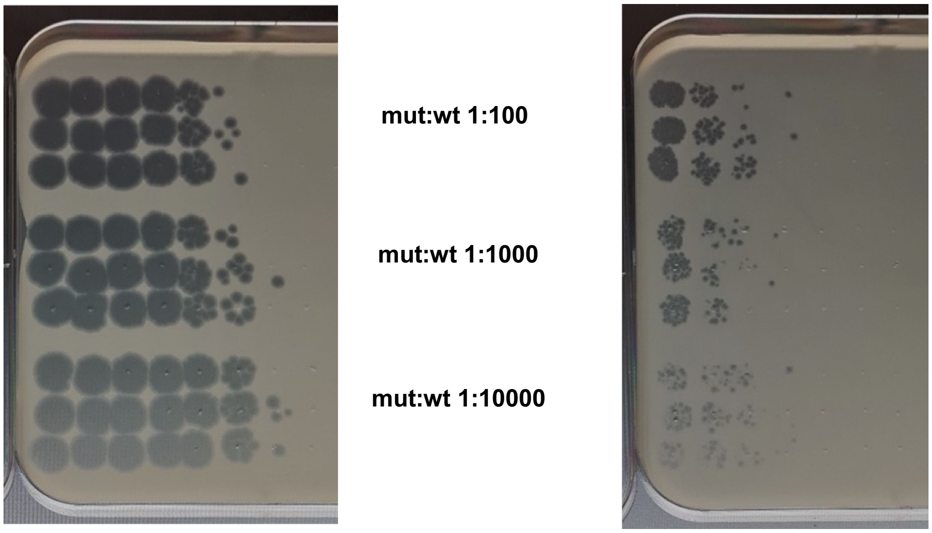

**Titers on *E. coli* B:**
mut:wt 1:100 $5.0 \pm 1.4 \times 10^{8}$ PFU/ml
mut:wt 1:1000 $1.5 \pm 0.3 \times 10^{9}$ PFU/ml
mut:wt 1:10000 $5.0 \pm 1.4 \times 10^{9}$ PFU/ml

**Titers on *rfaC*:**
mut:wt 1:100 $9.2 \pm 0.9 \times 10^{6}$ PFU/ml
mut:wt 1:1000 $4.1 \pm 2 \times 10^{6}$ PFU/ml
mut:wt 1:10000 $1.5 \pm 0.7 \times 10^{6}$ PFU/ml

**Supplementary Figure 17.** Dilution experiment: link between genotype and phenotype. **a.** Spotting assay of the tree first assemblies corresponding to T7D-S541R fragments dilution factor 0, 0.5, and 0.1 on strains *E. coli* B and *rfaC* at final 0.1 nM DNA concentration in TXTL.

**b.** Spotting assay of the tree last assemblies corresponding to T7D-S541R fragments dilution factor 0.01, 0.001, and 0.0001 on strains *E. coli* B and *rfaC* at final 0.1 nM DNA concentration in TXTL. The first spot corresponds to a dilution of 10. Each condition has three reactions spotted once.

**Supplementary Figure 18.** Evaluation of randomness of tail fiber assembly in TXTL. T7 contains six tail fibers. Each tail fiber is a trimer of gp17. In the case of the co-expression of the two-phage system in batch TXTL, we can make three hypotheses ordered by permissiveness: H0, H1, H2 on how T7 tail fibers are assembled in the absence of g/P coupling and pure mixing in TXTL. Here, pure (100%) g/P linked phage is defined as a phage that displays six tail fibers composed of only Mutant tail fibers if it encapsulates a mutant genotype or only WT tail fibers if it encapsulates a wt genotype. We further make the assumptions that Mutant and WT tail fibers are equally expressed and assemble at the same rates. Conversely, an inversely linked phage is defined as a phage that displays six tail fibers composed of only Mutant tail fibers if encapsulates a wt genotype or only WT tail fibers if it encapsulates a mutant genotype. We define D as the ratio of mutant genomes.

- In the equimolar co-assembly D is $\frac{1}{2}$
- In the dilution experiment D is $\frac{1}{2}$ ; $\frac{1}{10}$ ; $\frac{1}{100}$ ; $\frac{1}{1000}$ ; $\frac{1}{10000}$

**H0**: Hypothesis presented in the manuscript. 18 monomers assemble randomly in 6 tail fibers. If the phage has a mutant genotype, only 1 mutant monomer confers a ReLPS+ phenotype. In this condition:

(i) the fraction of pure g/P link and pure inverse g/P link in an equimolar mix is: $D^{18} = \frac{1}{2^{18}}$.

(ii) the fraction of phages that are ReLPS infectious phages is: $(1 - (1 - D)^{18}) \times D$

**H1**: Only identical monomers can assemble in tail fiber trimer. 6 tail fiber trimers assemble randomly onto a prophage. At least 1 mutant tail fiber confers ReLPS+ phenotype. In this condition:

(i) the fraction of pure g/P link and pure inverse g/P link in an equimolar mix is: $D^{6} = \frac{1}{2^{6}}$.

(ii) the fraction of phages that are ReLPS infectious phages is: $(1 - (1 - D)^{6}) \times D$

**H2**: 18 monomers assemble in 6 tail fiber trimers randomly. At least 1 tail fiber that is MMM confers ReLPS+ phenotype (MMM is a mutant tail fiber from 3 mutant monomers). In this condition:

(i) the fraction of pure g/P link and pure inverse g/P link in an equimolar mix is: $(D^{3})^{6} = \frac{1}{2^{18}}$.

(ii) the fraction of phages that are ReLPS infectious phages is: $(1 - (1 - D\ X\ D^{3})^{6}) \times D$

We compared the three hypotheses for the equimolar assembly experiment and the dilution experiment. Interestingly while H1 results in greatest fraction of pure link we still find experimentally find four times more pure linkage and the asymmetry between pure link and inversely link fractions. H1 however predicts less well than H0 for the dilution experiment when compared to the experimental results due to the condition that only identical monomers assemble in a tail fiber. Finally, H2 leads to the furthest predictions for both experiments. Indeed, Mutant monomers become quickly diluted in WT monomers reducing the chances that three identical monomers assemble in a tail fiber. Overall, these three simple hypotheses cover how tail fiber assemble in a perfectly mixed reaction in the absence of g/P coupling and do not predict our experimental results. While the equimolar co-expression experiment shows that bulk assembly results in only a small fraction of pure linkage, the g/P link is however not lost with dilution. This suggests that a coupling exist. This coupling could stem from an unknown interaction between phage proteins and its genome or the specific conditions of a TXTL reaction.

| Scenario | fraction | | | | |
|---|---|---|---|---|---|
| | wt/WT | wt/MIX | mut/MUT | mut/MIX | mut/WT & wt/MUT |
| Full linkage | 0.5 | 0 | 0.5 | 0 | 0 |
| H0 | $2 \times 10^{-6}$ | 0.499996 | $2 \times 10^{-6}$ | 0.499996 | $2 \times 10^{-6}$ |
| H1 | 0.0156 | 0.469 | 0.0156 | 0.469 | 0.0156 |
| H2 | $2 \times 10^{-6}$ | 0.499996 | $2 \times 10^{-6}$ | 0.499996 | $2 \times 10^{-6}$ |
| Experimental results | 0.06 ± 0.03 | 0.45 ± 16 | 0.06 ± 0.03 | 0.43 ± 16 | 0.0003 ± 0.0003 |

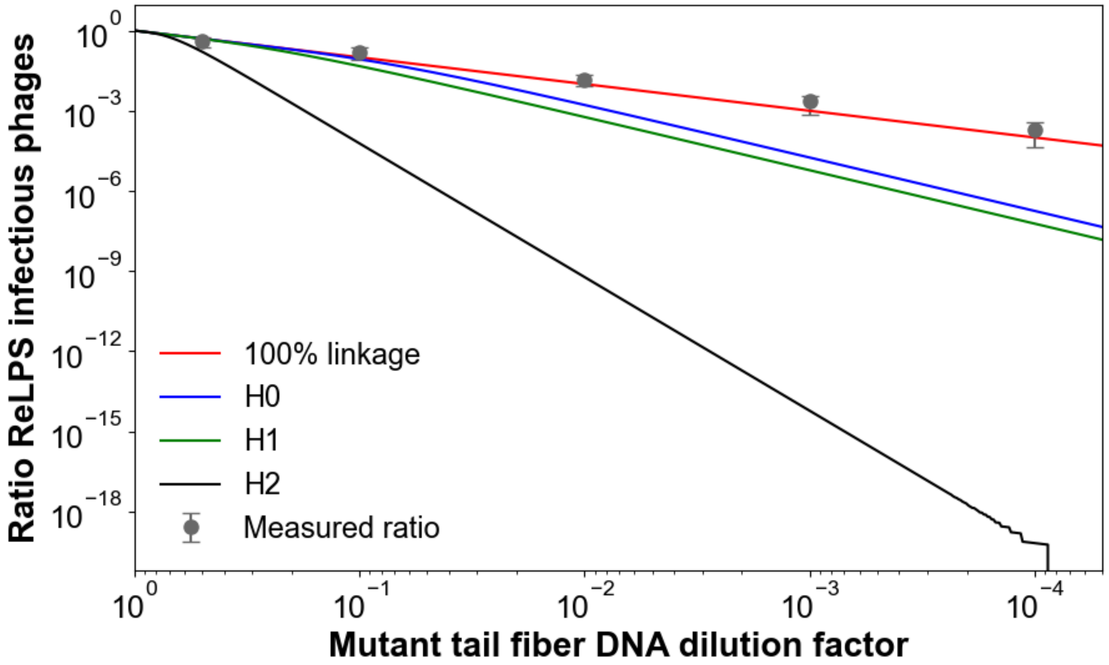

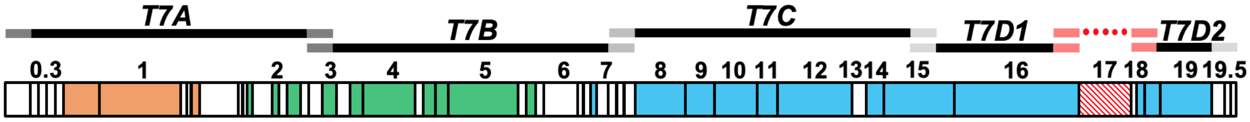

**E0**

| ref/mut | A | C | T | G |
|---|---|---|---|---|
| **A** | | 0.333 | 0.333 | 0.333 |
| **C** | 0.333 | | 0.333 | 0.333 |
| **T** | 0.333 | 0.333 | | 0.333 |
| **G** | 0.480 | 0.070 | 0.450 | |

**E1**

| ref/mut | A | C | T | G |
|---|---|---|---|---|
| **A** | | 0.333 | 0.333 | 0.333 |
| **C** | 0.333 | | 0.333 | 0.333 |
| **T** | 0.333 | 0.333 | | 0.333 |
| **G** | 0.607 | 0.117 | 0.276 | |

**E2**

| ref/mut | A | C | T | G |
|---|---|---|---|---|
| **A** | | 0.333 | 0.333 | 0.333 |
| **C** | 0.333 | | 0.333 | 0.333 |
| **T** | 0.333 | 0.333 | | 0.333 |
| **G** | 0.564 | 0.119 | 0.317 | |

**E3**

| ref/mut | A | C | T | G |
|---|---|---|---|---|
| **A** | | 0.333 | 0.333 | 0.333 |
| **C** | 0.333 | | 0.333 | 0.333 |
| **T** | 0.333 | 0.333 | | 0.333 |
| **G** | 0.538 | 0.121 | 0.341 | |

**T7-E0**

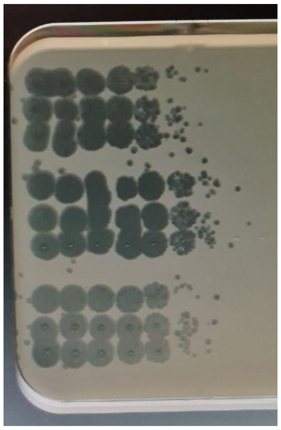

PFU/ml: $1.3 \times 10^{11}$
StDev: $1.1 \times 10^{11}$
EOP: 1

**T7-E1**

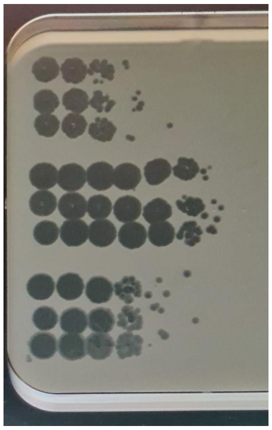

PFU/ml: $3.2 \times 10^{10}$
StDev: $5.5 \times 10^{10}$
EOP: 0.25

**T7-E2**

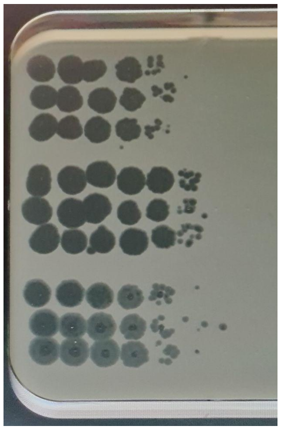

PFU/ml: $1.2 \times 10^{10}$
StDev: $1.4 \times 10^{10}$
EOP: 0.09

**T7-E3**

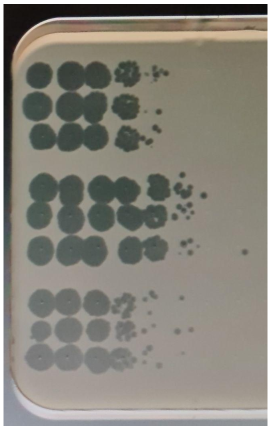

PFU/ml: $6.7 \times 10^{9}$
StDev: $9.8 \times 10^{9}$
EOP: 0.05

TXTL reaction + water

PHEIGES T7-E minus insert

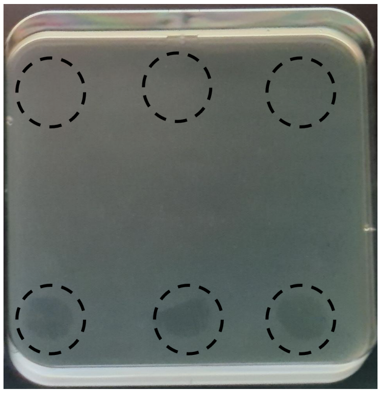

**Supplementary Figure 19.** Using PHEIGES to generate T7 phage variants with mutated tail fibers at four different rates of mutations. The genome was assembled from six DNA fragments. Four different mutated tail fiber tips DNA fragments (gene *17* 1146-1662) E0, E1, E2, E3 were obtained by PCR. The probability of nucleotide substitution for each fragment library are indicated in the tables. For instance, for E1, when a G is mutated, it is mutated to an A in 60.7% of the cases, to a C in 11.7% of the cases, and to a T in 27.6% of the cases (and the sum is 100%). Four T7 variants batches were spotted on an *E. coli* B lawn (T7-E0, T7-E1, T7-E2, T7-E3). The experiment was repeated three times, and each time spotted three times. The TXTL reaction was diluted in LB by factor of tens and spotted from left to right. The first spot on the left corresponds to a dilution of 100 of the TXTL reaction. Negative controls: (i) a TXTL reaction with added water (no DNA) was spotted three times on an *E. coli* B lawn, (ii) a TXTL reaction from PHEIGES without the insert (mutated tail fiber) but with all the other DNA parts was spotted three times on an *E. coli* B lawn without dilution. No plaques were formed. A slight inhibition of *E. coli* B growth is observed and explained later in the study **(see Fig. S28)**.

**Map of the spotting**

| | |
|---|---|
| T7-E0 | T7-E0 |
| T7-E1 | T7-E1 |
| T7-E1 | T7-E1 |
| T7-E2 | T7-E2 |
| T7-E3 | T7-E3 |
| T7-E0 | T7-E0 |
| T7-E1 | T7-E1 |
| T7-E1 | T7-E1 |
| T7-E2 | T7-E2 |
| T7-E3 | T7-E3 |

**Spotting plate**

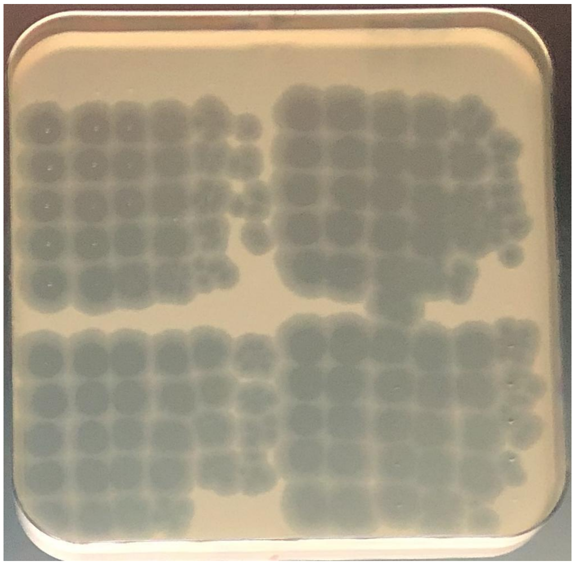

**Supplementary Figure 20.** Spotting assay of the four batches of T7 phages (T7-E0, T7-E1, T7-E2, T7-E3) on *E. coli* B strain. This figure shows the map of the spotting assay used in **Fig. S16** below. T7-E1 was spotted twice. Spotting was repeated four times on each plate. TXTL reactions from PHEIGES were diluted in LB by factor of ten and spotted from left to right. The first spot on the left corresponds to a dilution of 10 of the TXTL reaction.

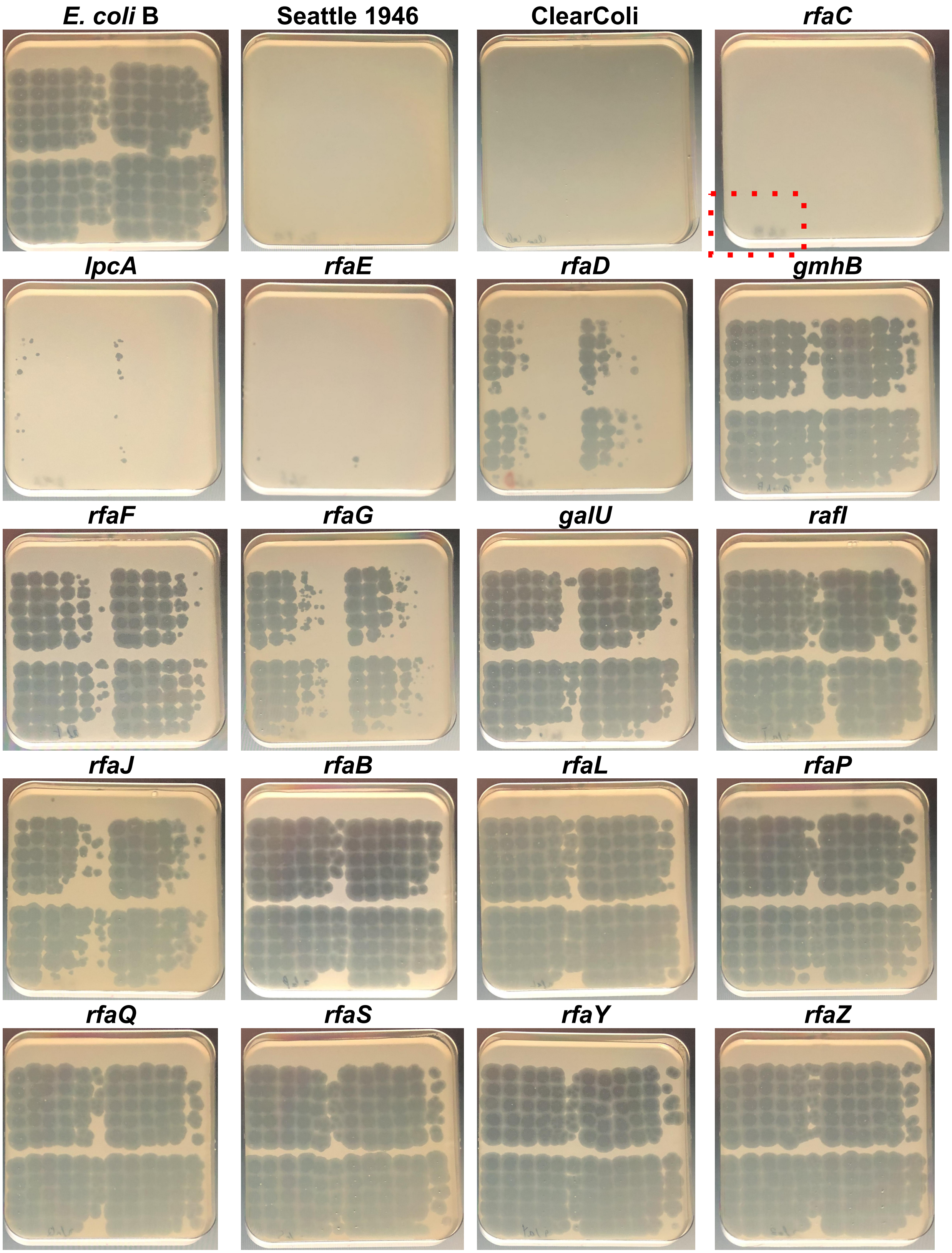
E. coli B
Seattle 1946
ClearColi
rfaC
lpcA
rfaE
rfaD
gmhB
rfaF
rfaG
galU
rafI
rfaJ
rfaB
rfaL
rfaP
rfaQ
rfaS
rfaY
rfaZ

| strains | EOP | | | |
|---|---|---|---|---|
| | **T7-E0** | **T7-E1** | **T7-E2** | **T7-E3** |
| *E. coli* B | 1 | 0.8 | 0.4 | 0.05 |
| Strain Seattle 1946 | no plaque | no plaque | no plaque | no plaque |
| ClearColi BL21(DE3) | no plaque | no plaque | no plaque | no plaque |
| *rfaC* | no plaque | no plaque | 2 plaques | no plaque |
| *lpcA* | no plaque | 14 plaques | 15 plaques | no plaque |
| *rfaE* | no plaque | 3 plaques | 5 plaques | no plaque |
| *rfaD* | 0.0004 | 0.0002 | 0.0001 | 0.000003 |
| *gmhB* | 0.8 | 0.6 | 0.2 | 0.04 |
| *rfaF* | 0.4 | 0.2 | 0.1 | 0.01 |
| *rfaG* | 0.04 | 0.04 | 0.04 | 0.0003 |
| *galU* | 0.6 | 0.2 | 0.1 | 0.0009 |
| *rfaI* | 0.8 | 0.5 | 0.5 | 0.1 |
| *rfaJ* | 0.1 | 0.04 | 0.03 | 0.0004 |
| *rfaB* | 0.8 | 0.6 | 0.3 | 0.01 |
| *rfaL* | 0.8 | 0.8 | 0.4 | 0.1 |
| *rfaP* | 0.6 | 0.5 | 0.3 | 0.0 |
| *rfaQ* | 0.6 | 0.5 | 0.3 | 0.1 |
| *rfaS* | 0.9 | 0.6 | 0.4 | 0.1 |
| *rfaY* | 0.8 | 0.6 | 0.5 | 0.1 |
| *rfaZ* | 0.9 | 0.9 | 0.4 | 0.2 |

**Supplementary Figure 21.** Spotting assay of the four batches of T7 phages (T7-E0, T7-E1, T7-E2, T7-E3) on all the *E. coli* strains listed in **Fig. S11**. **Fig. S15** shows the map. The red frame shown for *rfaC* indicates the location of plaques. The table shows the EOP.

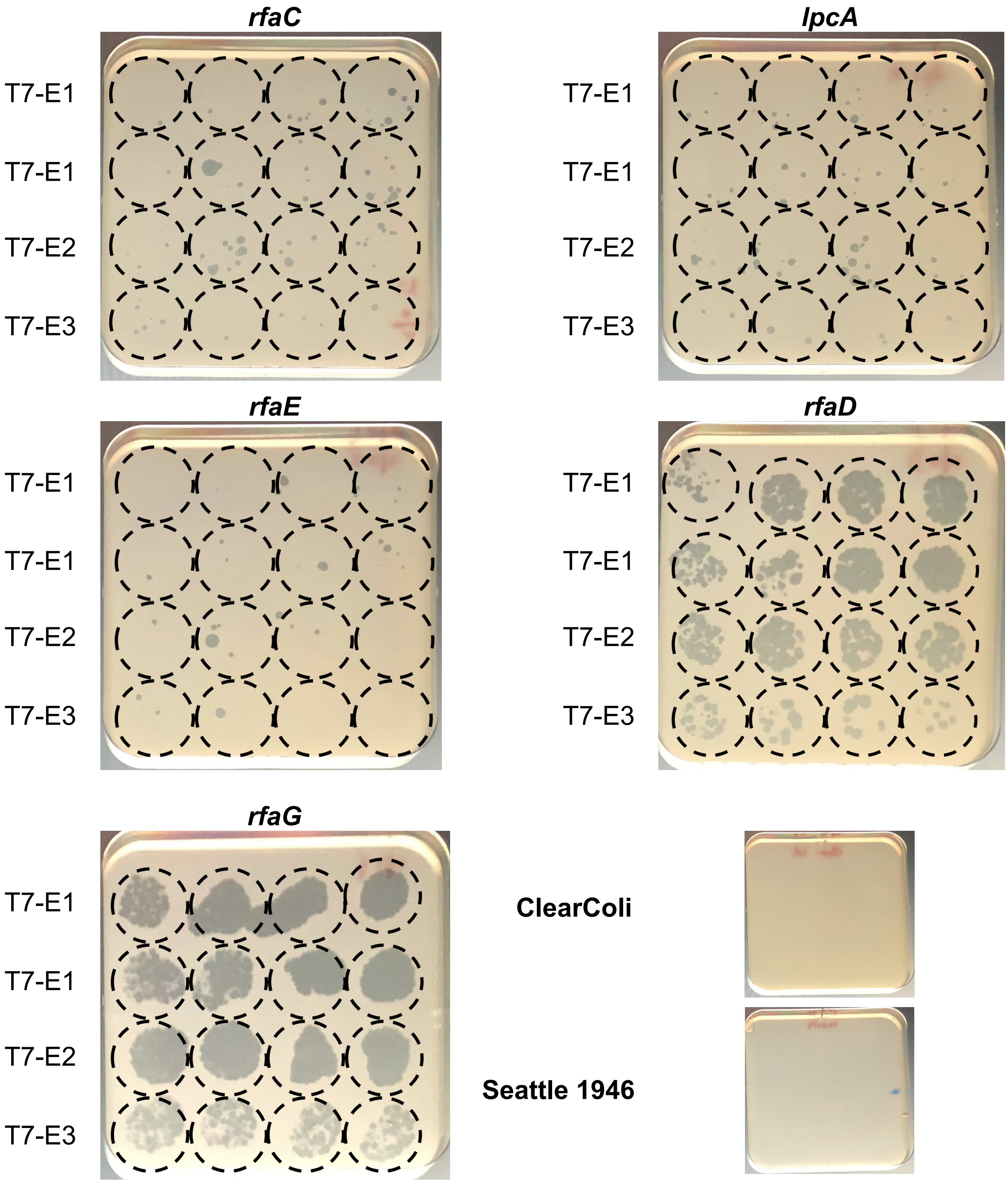

**Supplementary Figure 22.** Spotting of T7-E1, T7-E2 and T7-E3 on *rfaC*, *lpcA*, *rfaE*, *rfaD*, *rfaG*, ClearColi and Seattle 1946. No variant infected ClearColi and Seattle 1946. After PHEIGES, 10 µL of TXTL reaction were diluted hundred times. 25 µL of each dilution was spotted eight times for T7-E1 and four times for T7-E2 and T7-E3. A few to a dozen of plaques were detected on each ReLPS strain. Variants were picked for sequencing.

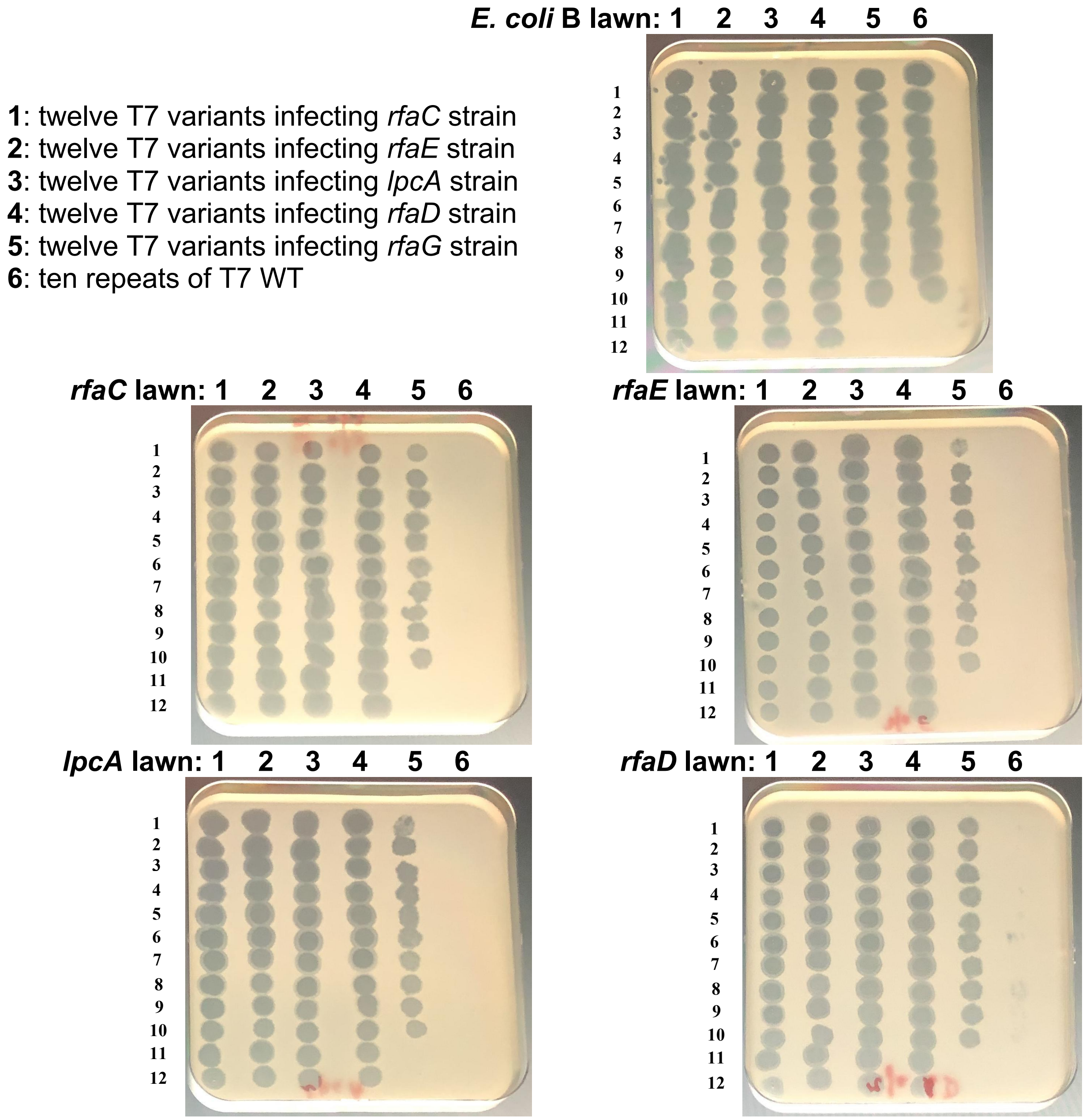

**Supplementary Figure 23.** Spotting assay to test T7 phage variants generated by PHEIGES on all the ReLPS *E. coli* strains (*E. coli* B used as a control, *rfaC*, *rfaE*, *lpcA* and *rfaD* mutant strains are all ReLPS). Each column 1-5 has twelve variants, column 6 is T7 WT. 6 first variants from T7-E1 (6 first from the top in Fig. 5d, labeled 1 to 6), 3 variants from T7-E2 (three following in Fig. 5d, labeled 7 to 9), and 3 variants from T7-E3 (last three in Fig. 5d, labeled 10 to 12). T7 WT phages were isolated from single plaques from T7-E0 assembly spotted on *E. coli* B. All the isolated variants amplified on their host could infect all the ReLPS strains except for some *rfaG* variants. No WT T7 infected ReLPS strains except *rfaD*. 3.5 µL of each undiluted clarified phage lysate were spotted.

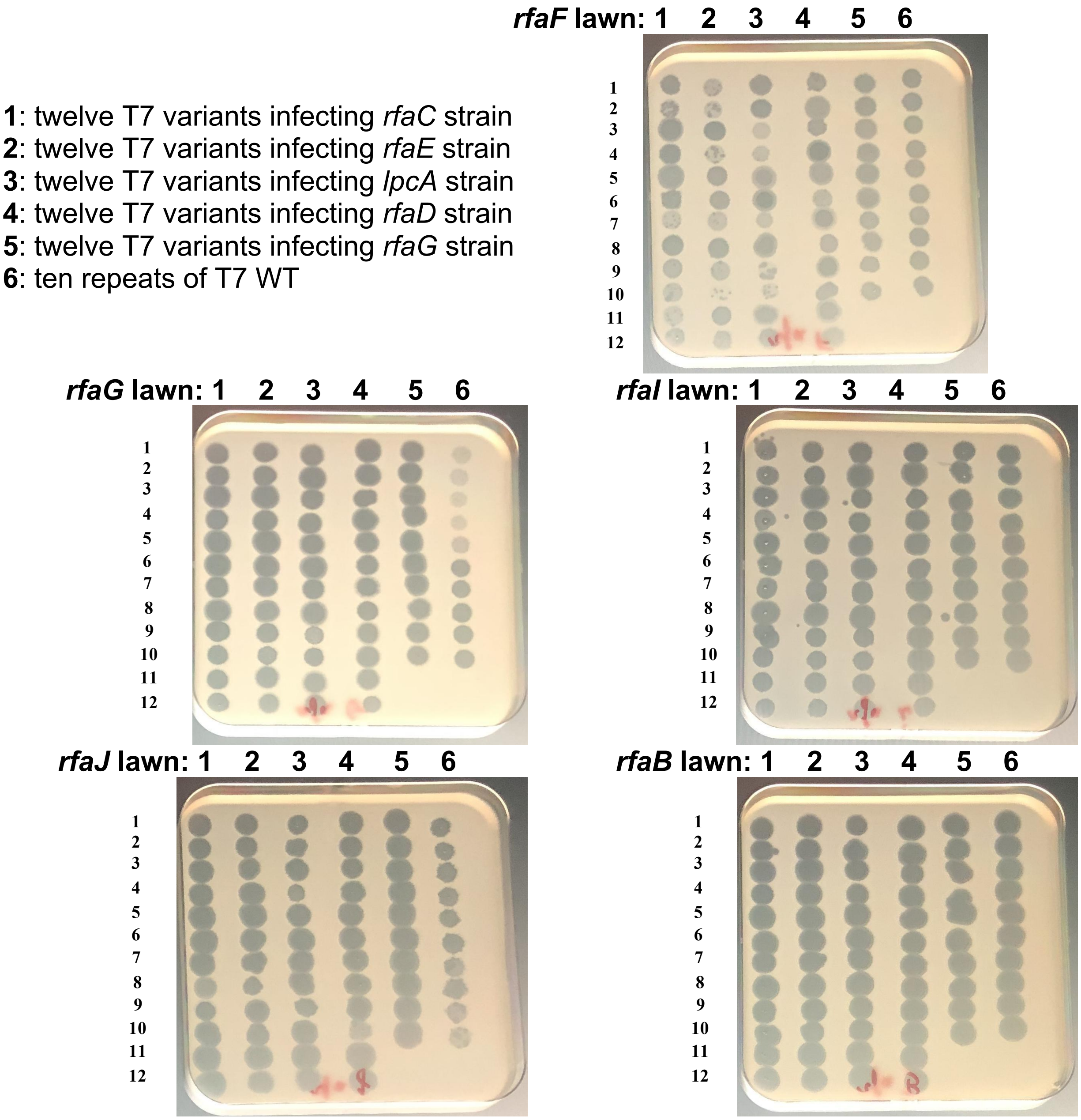

**Supplementary Figure 24.** Spotting assay to test T7 phage variants generated by PHEIGES on strains with different LPS: *rfaF* (Rd2LPS), *rfaG* (Rd1LPS), *rfaI* (RcLPS), *rfaJ* (RbLPS), *rfaB* (side chain modification LPS). Each column 1-5 has twelve variants (the same as in **Fig. S23)**, 6 first variants from T7-E1 (6 first from the top in Fig. 5d, labeled 1 to 6), 3 variants from T7-E2 (three following in Fig. 5d, labeled 7 to 9), and 3 variants from T7-E3 (last three in Fig. 5d, labeled 10 to 12), column 6 is T7 WT. T7 WT phages were isolated from single plaques from T7-E0 assembly spotted on *E. coli* B. Contraction of host range is visible on *rfaF*: some *rfaC*, *rfaE* and *lpcA* T7 variants were not able to infect the *rfaF* strain while T7 WT can. 3.5 µL of each undiluted clarified lysate were spotted.

### *rfaD*

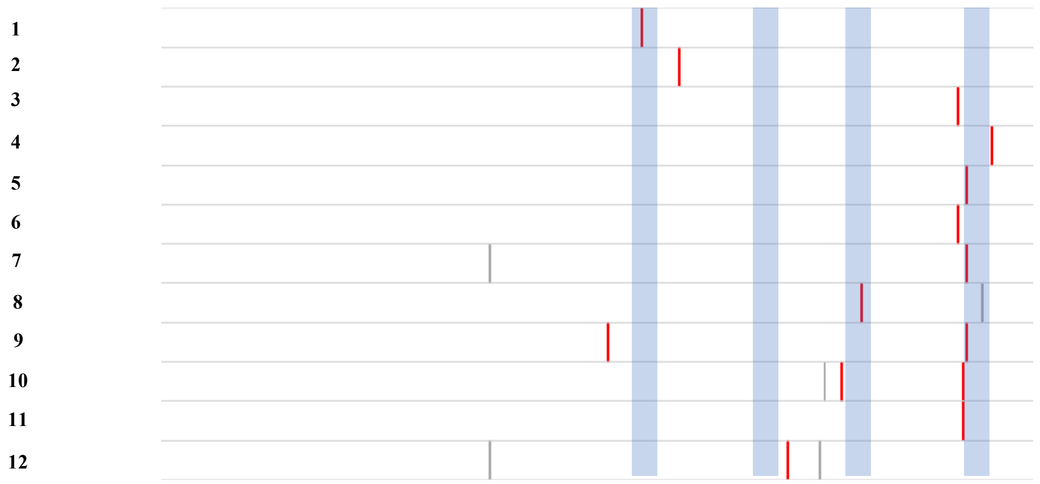

| | | | | | | | | |
|---|---|---|---|---|---|---|---|---|
| | R | | | | | | | |
| | | A | | | | | | |
| | | | | | | Y | | |
| | | | | | | | | K |
| | | | | | | | R | |
| | | | | | | N | | |
| | | | | | | | R | |
| | | | | | R | | | |
| S | | | | | | | R | |
| | | | | L | | | R | |
| | | | | | | | R | |
| | | | S | | | | | |
| T471 | S477 | T485 | F506 | I517 | G521 | D540 | S541 | N546 |

***rfaG***

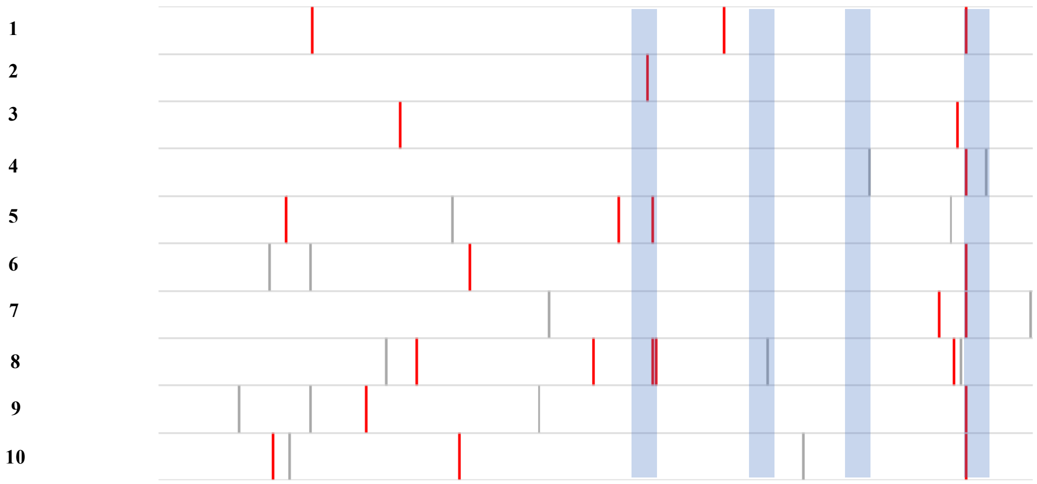

| | | | | | | | | | | | | | | | | |
|---|---|---|---|---|---|---|---|---|---|---|---|---|---|---|---|---|
| | | H | | | | | | | | | | H | | | | R |
| | | | | | | | | | | R | | | | | | |
| | | | | R | | | | | | | | | | | Y | |
| | | | | | | | | | | | | | | | | R |
| | S | | | | | | | | R | | R | | | | | |
| | | | | | | | D | | | | | | | | | R |
| | | | | | | | | | | | | | M | | | R |
| | | | | | Q | | | M | | | R | | | V | | |
| | | | H | | | | | | | | | | | | | R |
| I | | | | | | N | | | | | | | | | | R |
| N405 | C408 | Y413 | Q423 | K430 | H433 | D442 | N444 | K468 | V473 | G479 | G480 | N494 | K536 | A539 | D540 | S541 |

**Supplementary Figure 25.** Mutation landscape for twelve T7 phage variants generated by PHEIGES that infect *E. coli* mutant strains *rfaD* (ReLPS), 6 first variants from T7-E1 (6 first from the top in Fig. 5d, labeled 1 to 6), 3 variants from T7-E2 (three following in Fig. 5d, labeled 7 to 9), and 3 variants from T7-E3 (last three in Fig. 5d, labeled 10 to 12). and ten T7 variants that infect *rfaG* (ReLPS, all the variants are from T7-E3, labeled 1 to 10). These variants poorly infected by T7 WT. These mutation patterns are retrieved by ORACLE[5]. **Top row:** diagram showing the mutation landscape after treating the sequencing data. Silent mutations are grey and non-silent mutations are red. The blue zones show the external loops of the tail fiber tip. **Bottom row:** tables that show the annotated mutations. The bottom row in each table shows the positions of the native amino acids in the 172 amino acid segment (0 to 172, corresponding to amino acids 382 to 554 in gp17) that was mutated. The brackets show the amino acid substitution.

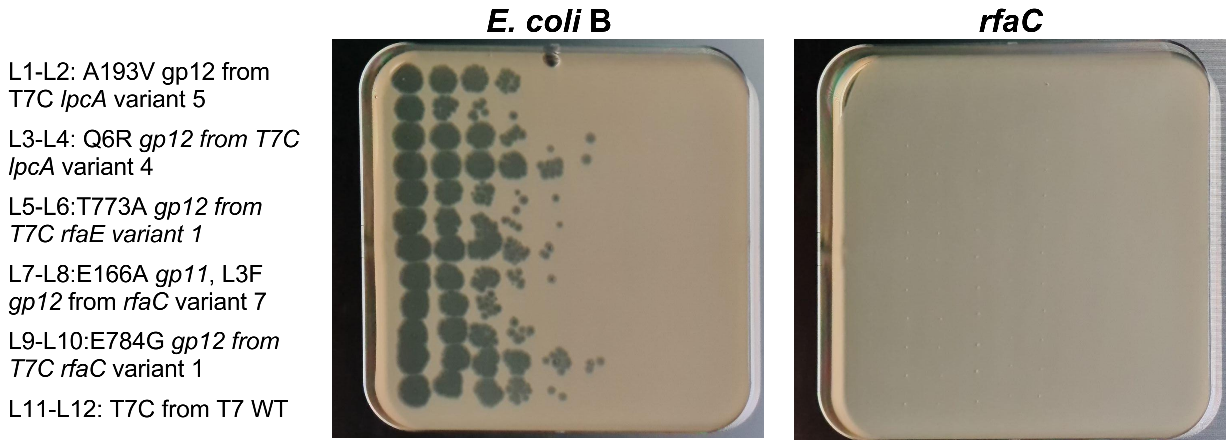

**Supplementary Figure 26.** PHEIGES was used to synthesize T7 phages with mutations only found in the tail genes *11* and *12*, by assembling genomes from four parts: T7A, T7B, T7D from T7 WT and T7C from variants as indicated in the figure. Each phage was assembled twice. Serial dilution was spotted once on *E. coli* B and *rfaC* mutant strain. No plaques on rfaC suggesting that the tail mutations are not responsible for the gain of function. The first spot on the left corresponds to a dilution of 10 of the TXTL reactions. A single plaque was picked and amplified in *E. coli* B for tail only lpcA variant 4 and rfaC variant 1. These phages were fully sequenced **(Table S3)**.

L1-L2: S541R gp17 from T7D *rfaC* 1

L3-L4: D540Y, G480S, S562G, G521R from T7D *rfaC* variant 7

L5-L6: S547R gp17 from T7D *rfaE* variant 9

L7-L8: G521R gp17 from T7D *lpcA* variant 4

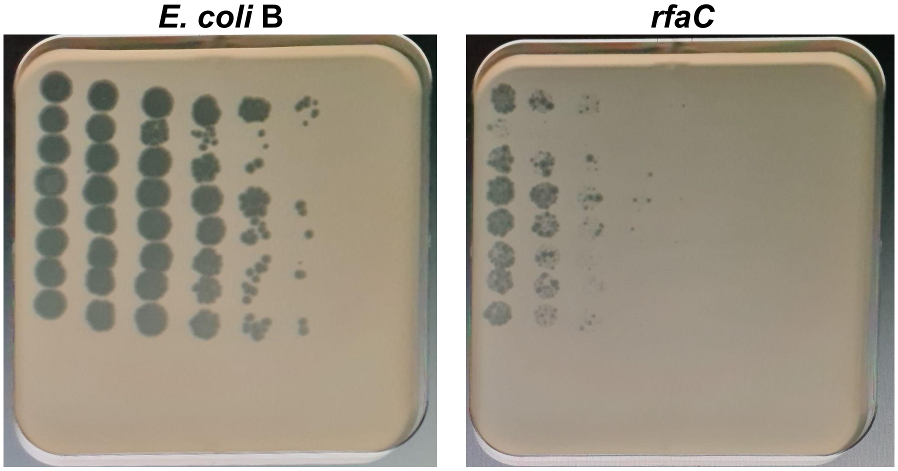

**Supplementary Figure 27.** PHEIGES was used to synthesize T7 phages with mutations only found in the tail fiber gene *17*, by assembling genomes from four parts: T7A, T7B, T7C from T7 WT and T7D from variants as indicated in the figure. Each phage was assembled twice. Serial dilution was spotted once on *E. coli* B and *rfaC* mutant strain. Low EOP are observed for the tail-fiber only assemblies on rfaC compared to *E. coli* B suggesting a poor fitness of the phage in absence of tail mutation in gene *11* and gene *12*. The first spot on the left corresponds to a dilution of 10 of the TXTL reactions. A single plaque was picked and amplifies in *E. coli* B for tail fiber only lpcA variant 4 and rfaC variant 1. These phages were fully sequenced **(Table S3)**.

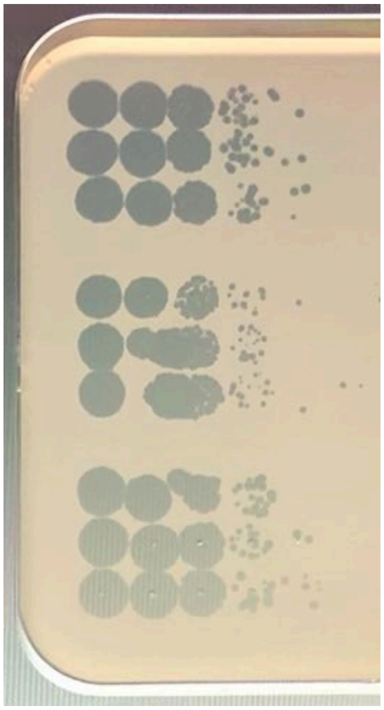

T7WT

T7gp10-3xFLAG

T7gp17-3xFlag

**Supplementary Figure 28.** Three PHEIGES assembly of T7WT, T7gp10-3xFLAG and T7gp17-3xFLAG. Tenfold dilutions of the TXTL reactions were spotted on a lawn of *E. coli* B. The first spot on the left corresponds to a dilution of 100 of the TXTL reaction. 3x-FLAG tags were fused to the C terminal of gp10B and gp17 by adding the 3x-FLAG sequence in primer overhangs. Primer sequences and assembly method are detailed in Supplementary Table S2. A single plaque for T7gp10-3xFLAG and T7gp17-3xFLAG were picked and their genome was sequenced to verify the presence of 3x-FLAG tag fusion (Sup. Table S1).

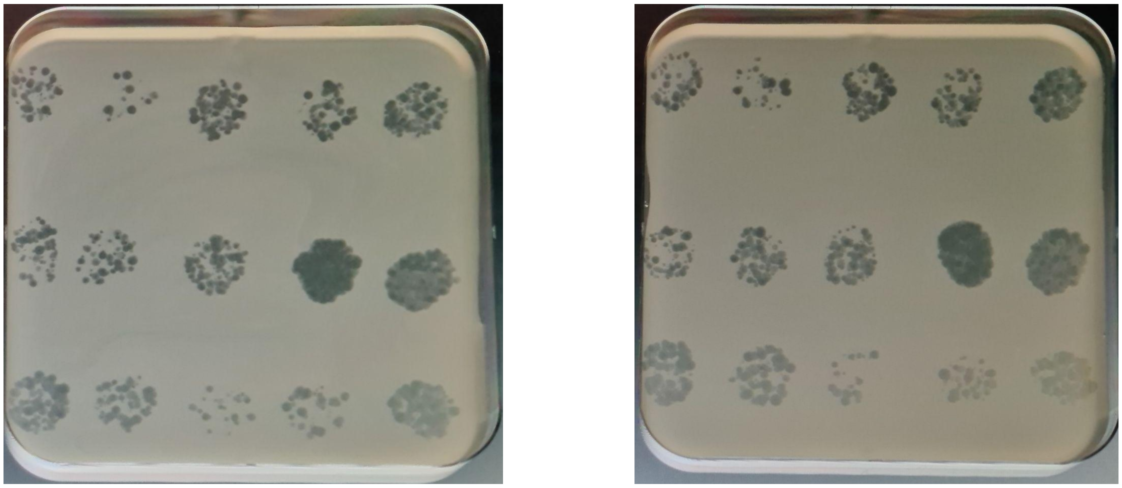

**Supplementary Figure 29.** Spotting of the engineered T7-compilation phage obtained using PHEIGES, on a lawn of the *E. coli rfaC* mutant strain. Fifteen assemblies were made and spotted twice (left and right). Each assembly included a different ReLPS T7D part from the collection of T7 variant phages that infect *E. coli* mutant strains with ReLPS (five from the *rfaC* mutant strain, five from the *lpcA* mutant strain and five from the *rfaE* mutant strain). All yielded phages on the *rfaC* mutant strain indicating ReLPS mutant. The TXTL reaction was diluted 100 times and 25 µL of this dilution were spotted twice. Three T7-compile phages with S541R mutation in gp17 were picked, amplified in *E. coli* B, and fully sequenced. The three genomes contained the deletion and mCherry cassette insertion as well as different mutation in tail genes *11* and *12*.

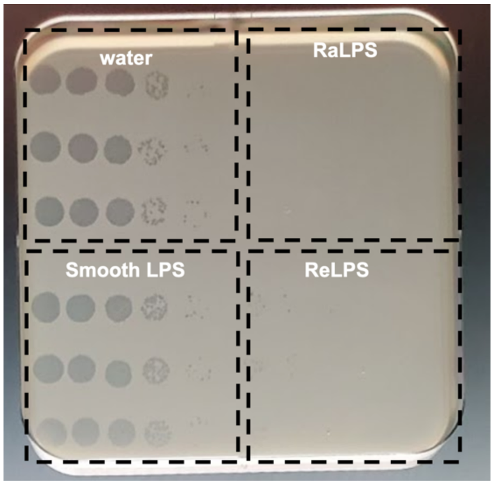

T7-compilation phage spotting on a lawn of *E. coli* B after incubation with:

| | mean PFU/ml | StDev |
|---|---|---|
| + water | $1.5 \times 10^8$ | $5.0 \times 10^7$ |
| + smooth LPS | $1.6 \times 10^8$ | $5.2 \times 10^7$ |
| + RaLPS | $< 10^3$ | - |
| + ReLPS | $\sim 10^4$ | - |

**Supplementary Figure 30.** In vitro genome ejection assay using purified LPS. Spotting (on a lawn of *E. coli* B) of serial 10-fold dilutions of the T7-compilation phage generated by PHEIGES. The phage at $10^8$ PFU/mL was pre-incubated with purified LPS at 0.2 mg/mL for 2 h at 37 °C (mild conditions). RaLPS is the full size LPS. In the case of the T7-compilation, both the RaLPS and the ReLPS induce ejection of the genome during the pre-incubation resulting in no plaques for RaLPS and a few undefined plaques on the first spot for ReLPS. The first spot on the left corresponds to no dilution.

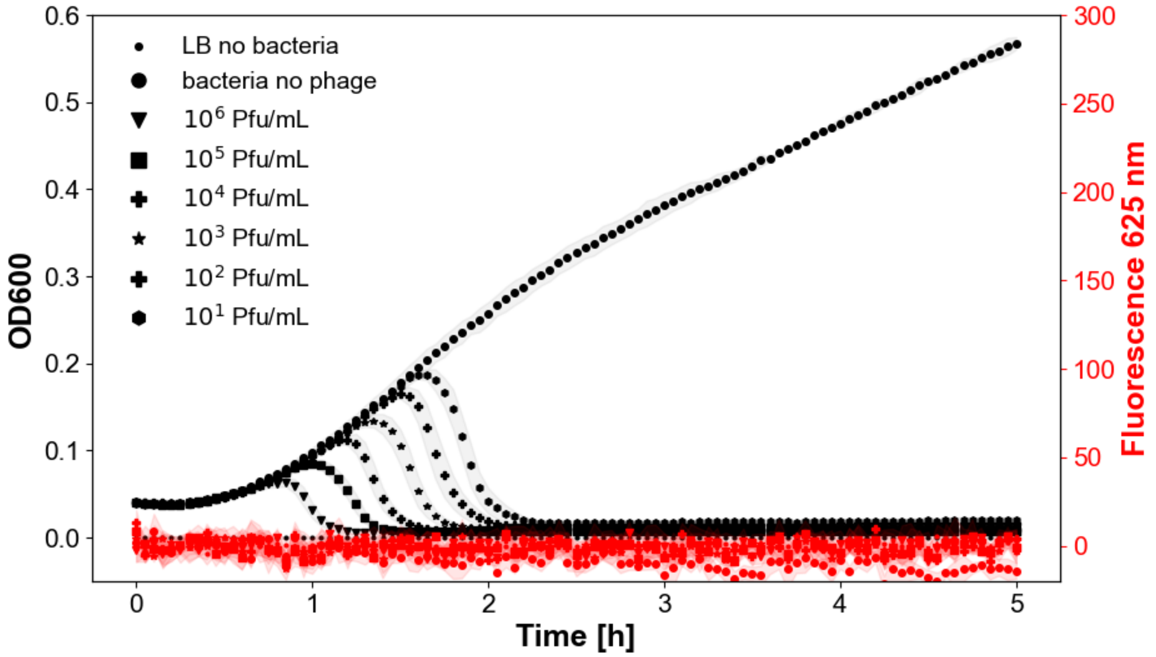

| dilution | $10^1$ | $10^2$ | $10^3$ | $10^4$ | $10^5$ | $10^6$ |
|---|---|---|---|---|---|---|
| **Time ratio** $\frac{Tcomp - Tmut}{Tcomp}$ | 0.19 | 0.259 | 0.27 | 0.307 | 0.33 | 0.35 |

**Supplementary Figure 31.** Kinetics of infection of *E. coli* mutant strain *rfaC* by a T7-ReLPS phage. Left: OD600 (dark curves). Right: fluorescence at 625 nm (red curves). T7-compilation and Tmut corresponds to the time at which the OD600 is maximal at a given dilution factor for T7-compilation and T7-ReLPS. Compared to the T7-compilation **(Fig. 5b)**, at the same phage concentrations the T7-ReLPS is on average ~30% faster at clarifying the rfaC cultures.

Phage T4
169 kbp
GenBank: NC_000866
Plaque assay on a lawn of *E. coli* B
~$10^8$ PFU/ml

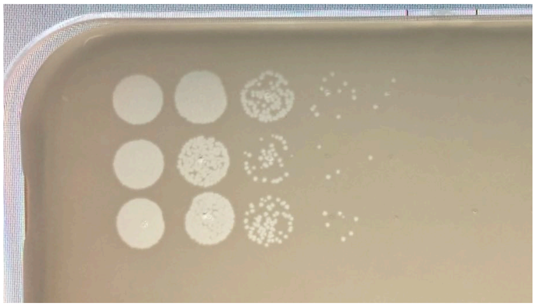

Phage T6
169 kbp
GenBank: NC_054907
Spotting assay on a lawn of *E. coli* B
~$10^8$ PFU/ml

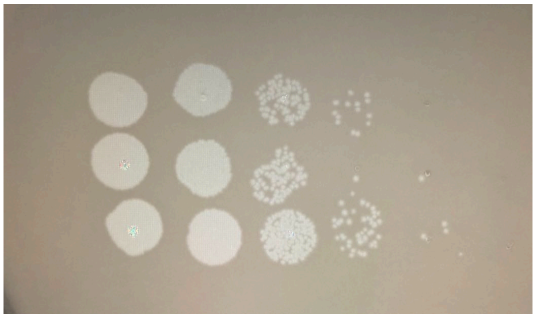

Phage VpaE1
88 kbp
GenBank: NC_027337.1
Plaque assay on a lawn of *E. coli* $B^E$
~$10^{10}$ PFU/ml

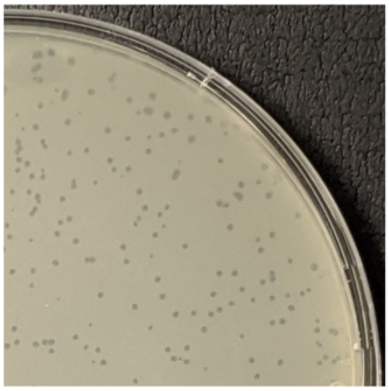

Phage FelixO1
86 kbp
GenBank: NC_005282
Plaque assay on a lawn of *Salmonella* LT2
~$10^8$ PFU/ml

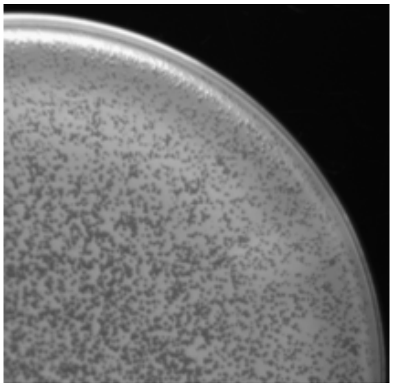

Phage S16
160 kbp
GenBank: NC_020416
Plaque assay on a lawn of *Salmonella* LT2
~$10^8$ PFU/ml

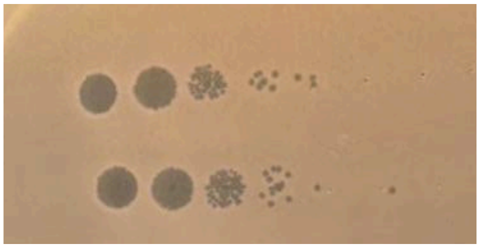

**Supplementary Figure 32.** TXTL expression of various phages from their purified genomes at 0.1nM. **Phage T4 and T6:** Three independent expression of T4 genome at 0.1nM in TXTL, plaque assay on *E. coli* B. **VpaE1**: plaque assay of the *E. coli* phage VpaE1[6] on a lawn of *E. coli* $B^E$ [6]. VpaE1 was provided by Lidija Truncaite, Vilnius University, Lithuania. **FelixO1 and S16:** plaque assay of the *Salmonella* phage FelixO1[7] and S16[8] on a lawn of *Salmonella* LT2. FelixO1 and S16 were provided by Kenneth Sanderson at the University of Calgary, Salmonella Genetic Stock Center. The five genomes were extracted and purified as reported before[9]. The TXTL reactions and plaque assays were performed as for phage T7. No plaques were observed when the genomic DNA of the five phages was spotted.

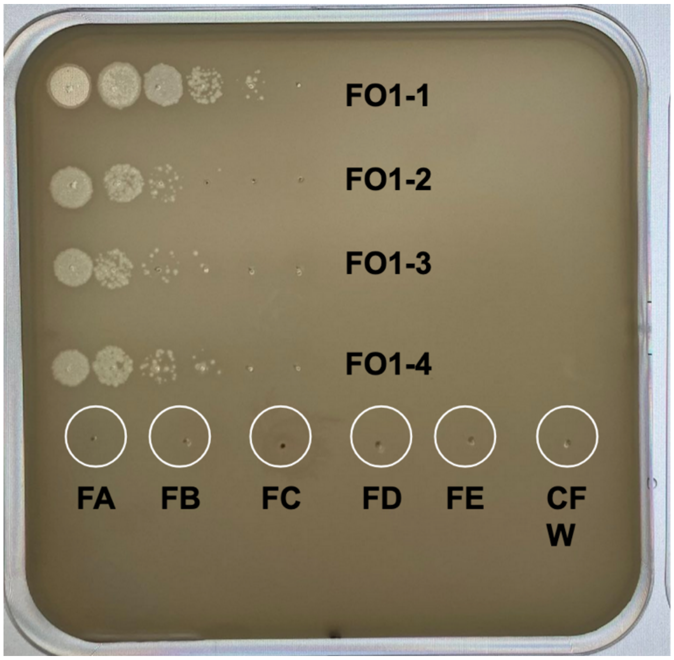

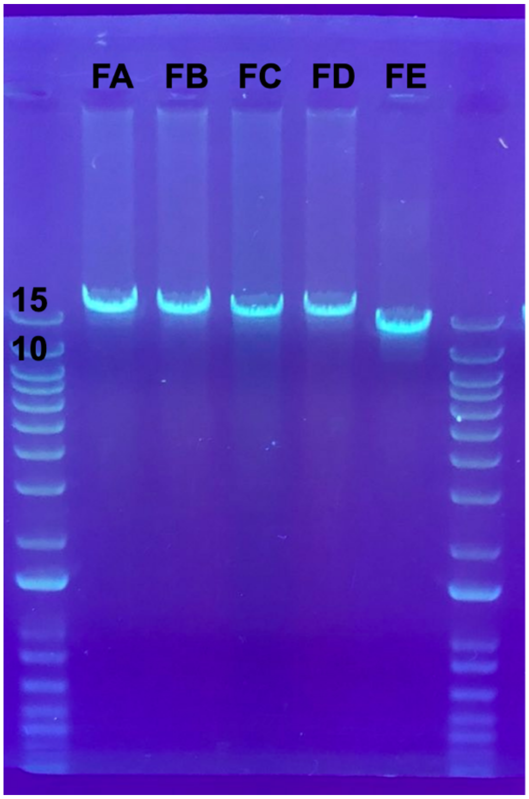

**Supplementary Figure 33.** FelixO1 assembly from 5 fragment (FA 18709 bp, FB 17504 bp, FC 18838 bp, FD 18409 bp, FE 13045 bp). 4 independent replicate assembly 0.1nM final (FO1-1, FO1-2, FO1-3, FO1-4) yielding 7.2 +/- 0.5 x$10^7$ PFU/mL. Tenfold TXTL reaction serial dilutions in LB spotted once each (3.5uL). The first spot is dilution 10. Negative controls are each FelixO1 fragment expressed in cell-free and cell-free water, spotted once 10uL. Left is DNA gel electrophoresis, ladder is 1kb+. PCR fragments sequences and primers are detailed in Table S1 and S2.

#### References

1. Garamella, J., Marshall, R., Rustad, M. & Noireaux, V. The All E. coli TX-TL Toolbox 2.0: A Platform for Cell-Free Synthetic Biology. *ACS Synth Biol* **5**, 344–355 (2016).
2. Garenne, D., Thompson, S., Brisson, A., Khakimzhan, A. & Noireaux, V. The all-E. coliTXTL toolbox 3.0: new capabilities of a cell-free synthetic biology platform. *Synthetic Biology* **6**, ysab017 (2021).
3. Häuser, R. *et al.* Bacteriophage protein-protein interactions. *Adv Virus Res* **83**, 219–298 (2012).
4. Washizaki, A., Yonesaki, T. & Otsuka, Y. Characterization of the interactions between Escherichia coli receptors, LPS and OmpC, and bacteriophage T4 long tail fibers. *Microbiologyopen* **5**, 1003–1015 (2016).
5. Huss, P., Meger, A., Leander, M., Nishikawa, K. & Raman, S. Mapping the functional landscape of the receptor binding domain of T7 bacteriophage by deep mutational scanning. *Elife* **10**, e63775 (2021).
6. Šimoliūnas, E. *et al.* Incomplete LPS Core-Specific Felix01-Like Virus vB_EcoM_VpaE1. *Viruses* **7**, 6163–6181 (2015).
7. Whichard, J. M. *et al.* Complete genomic sequence of bacteriophage felix o1. *Viruses* **2**, 710–730 (2010).
8. Dunne, M. *et al.* Salmonella Phage S16 Tail Fiber Adhesin Features a Rare Polyglycine Rich Domain for Host Recognition. *Structure* **26**, 1573-1582.e4 (2018).
9. Rustad, M., Eastlund, A., Marshall, R., Jardine, P. & Noireaux, V. Synthesis of Infectious Bacteriophages in an E. coli-based Cell-free Expression System. *J Vis Exp* 56144 (2017) doi:10.3791/56144.
